# The epidermal stem cell-supporting matricellular protein fibulin 7 modulates skin inflammatory response in a psoriasis model

**DOI:** 10.64898/2026.03.31.715486

**Authors:** Erna Raja, Tomoko Machida, Narenmandula, Karolina Edlund, ASM Sakhawat Hossain, Fan Wenxin, Jun Tsunezumi, Yukihide Watanabe, Keiichi Asano, Kenichi Kimura, Ken Natsuga, Aiko Sada, Hiromi Yanagisawa

## Abstract

ECM composition and organization are greatly altered during inflammation but it is still elusive if ECM dynamics may protect tissue stem cells against aberrant inflammation. Fibulin 7 (encoded by *Fbln7*) is part of the basement membrane ECM where epidermal stem cells (EpSCs) reside. It supports the long-term potential of fast-cycling EpSCs and moderates aging-related inflammatory markers in keratinocytes. Here, we assessed fibulin 7’s role during imiquimod (IMQ)-induced inflammation in 1-year-old mouse dorsal skin. We found that loss of *Fbln7* aggravates epidermal inflammation, marked by increased epidermal thickness, proliferation, and phosphorylation of JNK (c-Jun N-terminal kinase). Fast-cycling EpSCs labeled with *Slc1a3*-creER-TdTomato demonstrated that IMQ-induced proliferation in *Fbln7* KO mice is contributed by cell divisions in the suprabasal layers, a hallmark of inflammatory epidermal responses. EpSC transcriptomes further reveal IMQ-modulated genes that are more substantially affected in *Fbln7* KO mice, including IL-17 pathway-related genes known in psoriasis pathogenesis. Mechanistically, fibulin 7 directly binds to IL-17A and decreases IL-17A-mediated p38 MAPK activation. In public human psoriasis datasets, *FBLN7* is reduced in lesional skin compared with non-lesional or normal skin, and it is significantly correlated with common psoriasis-associated genes. Altogether, fibulin 7 is potentially beneficial to protect against skin inflammation.

## Introduction

The ECM is a crucial component of the tissue microenvironment, providing not only cell adhesion sites but also regulating cell behavior by modulating the bioavailability of growth factors/cytokines, binding to cell receptors for signaling and mechanotransduction (Gattazzo et al., 2014). In the skin, ECM homeostasis is maintained by a balance between ECM secretion and degradation mediated by fibroblast- and keratinocyte-derived enzymes (Pfisterer et al., 2021). Similarly, EpSCs within basal keratinocytes produce and organize their own niche, including BM proteins and various growth factors, which in turn maintain the balance between self-renewal and differentiation. EpSCs committed for differentiation leave the basal layer, forming the upper suprabasal layers and stratified epithelium (Fuchs and Blau, 2020).

When inflammation occurs, however, the influx of immune cells and cytokine-mediated signaling remodels the EpSCs niche via secretion of proteases such as matrix metalloproteases (MMPs) and elastases that generate bioactive ECM fragments, and various cells express matricellular proteins that support inflammatory processes (e.g. tenascins, periostin, CCN) (Bhattacharjee et al., 2019, Sorokin, 2010, Sutherland et al., 2023). During inflammatory stress, EpSCs respond by undergoing proliferation or cell death, lineage plasticity, inflammatory memory retention, and chemokines secretion to recruit immune cells (Xing and Naik, 2020). As a counterbalance, intact vasculature BM acts as a physical ‘barrier’ to prevent excessive immune cells infiltration, thereby controlling the extent of inflammation (Sutherland et al., 2023). However, much is unknown about whether epidermal BM ECM can shield EpSCs against detrimental effects of inflammation.

Psoriasis is a common recurring inflammatory disease characterized by thick, red, scaly skin. Systemically it is often associated with arthritis and cardiovascular co-morbidity risks (Gelfand, 2024). Psoriasis pathogenesis involves the classical immune hyperactivation along with other cell types amplifying the inflammatory response such as keratinocytes (Daccache and Naik, 2024, Ma et al., 2023, Shen et al., 2025, Wang and Lai, 2024, Zhou et al., 2022). Furthermore, psoriatic epidermis exhibits BM modifications, yet its biological relevance is still a discussion (Bozo et al., 2021, Flink et al., 2023, McFadden and Kimber, 2016, Natsumi et al., 2018).

Here we aim to assess whether fibulin 7, a stem cell-supporting matricellular protein in the epidermal BM, could limit keratinocyte hyperactivation in a mouse psoriasis model. Fibulin 7 has been reported to promote the longevity of fast-cycling EpSCs, and its loss of function was associated with upregulation of inflammation-related genes in aging EpSCs (Raja et al., 2022). Since ECM can bind cytokines and growth factors through glycosaminoglycans and proteoglycans (Gray et al., 2022), we hypothesized that fibulin 7 functions as a cytokine-binding ECM that regulates psoriasis development. In this study, *Fbln7* depletion resulted in a worsened phenotype in the dorsal skin of imiquimod (IMQ)-treated mice. In human skin, fibulin 7 protein co-localizes with Slc1a3-expressing basal cells at the rete ridges, which have been identified as fast-cycling EpSCs (Ghuwalewala et al., 2022, Sada et al., 2016). Fate analyses of these cells in IMQ-treated mouse skin combined with total EpSC transcriptome and in vitro assays suggested that fibulin 7 binds and modulates IL-17-mediated inflammatory response in EpSCs. Lastly, analysis of human datasets showed decreased *FBLN7* expression in psoriatic skin and an inverse relationship between *FBLN7* and psoriasis-related genes.

## Results

### Fibulin 7 suppresses epidermal hyperproliferation in an imiquimod-induced mouse skin inflammation model

IMQ is a known chemical ligand for Toll-like Receptor (TLR) 7 and its topical application on mouse dorsal skin induces inflammatory processes mimicking human psoriasis development (van der Fits et al., 2009). To investigate the role of fibulin 7 in IMQ-induced epidermal hyperplasia, IMQ cream was applied to *Fbln7*^+/+^ (WT) and *Fbln7*^-/-^(KO) mice in a short-term 4-day treatment (Fraillon et al., 2025) (Fig. 1A). Middle-aged 1-year-old (yo) mice were utilized due to previously reported aging features in 1-yo *Fbln7* KO mice (Raja et al., 2022) and to model late-onset psoriasis in an aging population (Ferrandiz et al., 2002, Goto et al., 2021).

**Figure 1.**
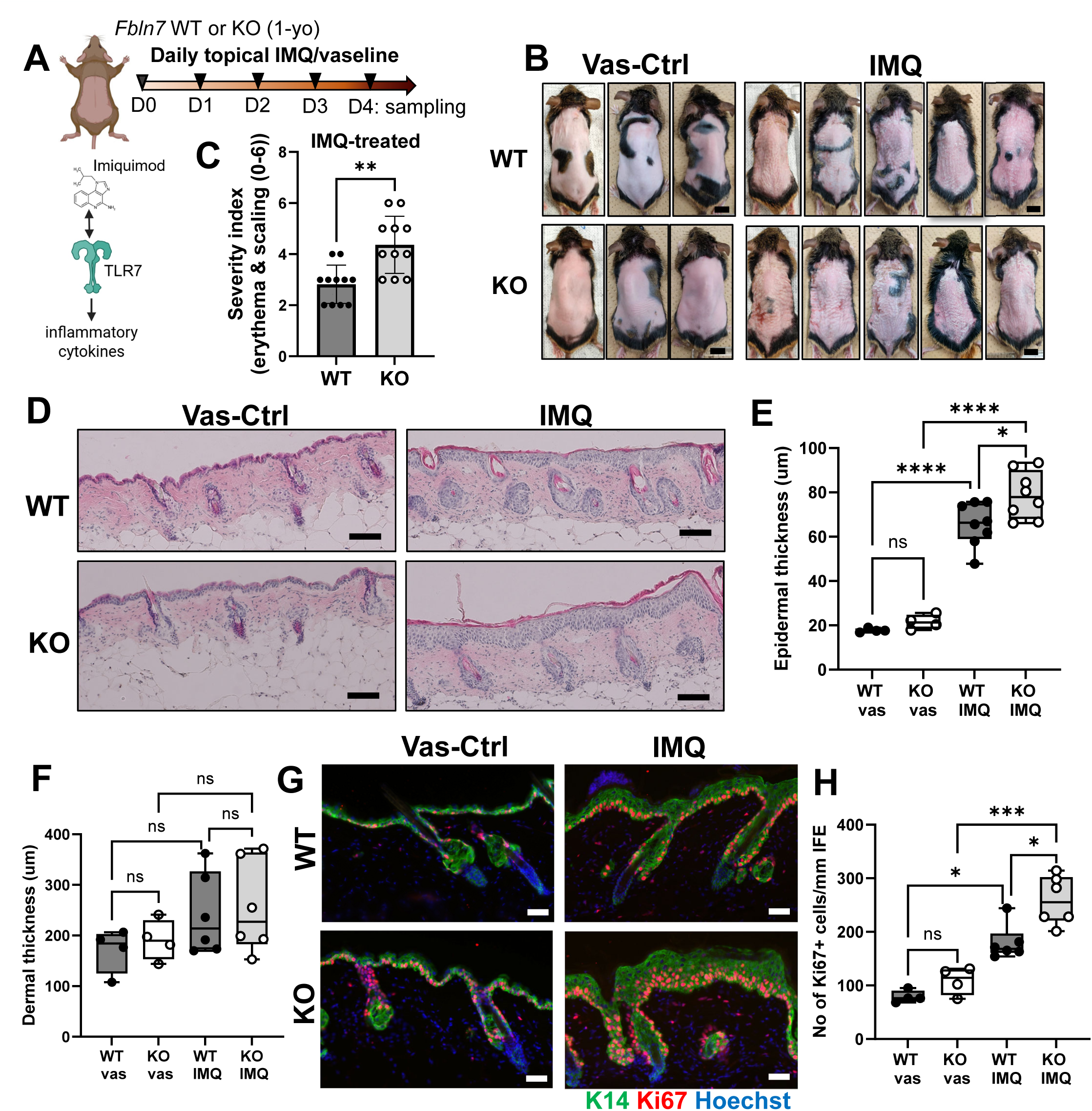
Loss of fibulin 7 augmented epidermal thickness and cell proliferation induced by imiquimod treatment on mouse dorsal skin. **(A)** Experimental timeline of IMQ/vas application for 4 times prior to skin sampling (D, day). IMQ activates toll-like receptor 7 (TLR7) initiating inflammatory cascades and further cytokines secretion. (B) Representative pictures of *Fbln7* WT or KO mice skin on sampling day 4. Ctrl, control. Scale bars, 1 cm. (C) Gross skin assessment on day 4, based on the extent of erythema (redness) and scaling (dryness). n = 11 per group, Welch’s unpaired T-test. (D) Hematoxylin and eosin (H&E) staining images of dorsal skin from each representative group at day 4 (Scale bars, 100 μm), and (E) epidermal thickness measured based on the H&E staining. vas-treated groups; n = 4, IMQ-treated groups; n = 8. ns, not significant; 2-way ANOVA, Fisher’s test. (F) H&E images was used to measure dermal thickness (area between epidermis and adipose tissue) after vas-control or IMQ treatment for 4 days. n = 4 for vas-treated groups and n = 6 for IMQ-treated groups. ns, not significant. 2-way ANOVA, Tukey test. (G) Immunofluorescence images of WT or KO skin stai*n*ed with anti-keratin 14 (K14) and Ki67 after treatment with vas or IMQ at day 4. Hoechst, nuclear counter stain. Scale bars, 50 μm. (H) Quantification of total Ki67-positive cells per mm interfollicular epidermis (IFE, epidermal areas excluding hair follicles), n = 4 for vas-treated groups and n = 6 for IMQ-treated groups. statistical test: 2-way ANOVA, Tukey test. * *P* < 0.05, ** *P* < 0.01, *** *P* <0.001, ns; not significant.

After IMQ application, dorsal skin became flaky dry with some irritation. Comparing between *Fbln7* WT and KO mice under IMQ induction, KO skin was more severely affected and appeared scalier with some increase in erythema (Fig. 1B, C). Histology indicated epidermal hyperplasia that was worse in KO compared to WT in IMQ-treated mice, despite no significant changes in dermal thickness (Fig.1D, E, F). Suprabasal keratin 14 further illustrated the epidermal inflammatory condition and a higher number of proliferating Ki67+ cells were observed in the KO skin (Fig. 1G, H) (Seldin and Macara, 2020). Thus, fibulin 7 weakens IMQ-induced epidermal hyperplasia.

### Slc1a3-expressing fast-cycling epidermal stem cell dynamics in the *Fbln7* KO mice after IMQ treatment

We previously reported that fast-cycling EpSCs express *Slc1a3* in the mouse skin and fibulin 7 participates in the maintenance of these stem cells (Raja et al., 2022, Sada et al., 2016). In the human skin, Slc1a3+ basal cells are located in the epidermal rete ridges and express markers resembling the fast-cycling EpSCs in mice (Ghuwalewala et al., 2022, Ishikawa et al., 2025). Fibulin 7 was found more abundant in the rete ridges BM underlying the Slc1a3+ basal cells, implying its relevance in supporting this stem cell population (Fig. S1A).

We then tested if Slc1a3+ EpSCs in the interfollicular epidermis (IFE/epidermal) changes their behavior in the absence of fibulin 7 during the short-term IMQ-induced inflammation (Fig. S1B). Using a previously established lineage tracing mouse model (*Slc1a3-*creER-tdTomato (*Slc1a3*-tdT+)) in 1-year-old *Fbln7* WT and KO mice, at which time the number of *Slc1a3*+ fast cycling cells was significantly decreased in *Fbln7* KO mice, fast-cycling EpSC responses to IMQ were analyzed. We observed that across all 4 conditions, a subset of cells remained in the basal layer, while others differentiated to the suprabasal layers (Fig. S1C). Consistent with our previous observation, *Slc1a3-*tdT+ basal epidermal cells were fewer in the *Fbln7* KO mice compared to WT (Fig. S2A – S2C). In contrast, IMQ treatment did not alter the number of basal epidermal *Slc1a3*-tdT+ cells (Fig. S2B, S2C).

To determine the proliferation status of these cells, Ki67+ cells were evaluated (Fig. S2D – S2G). The results suggest that after IMQ treatment, overall *Slc1a3-*tdT+ cells were more proliferative in the *Fbln7* KO epidermis compared to WT (Fig. S2E). However, the number of Ki67+ basal cells was not significantly increased under the influence of IMQ in *Fbln7* KO mice (Fig. S2F). Instead, the observed increase in proliferation marker was mainly contributed by *Slc1a3-*tdT+ cells in the suprabasal layers (Fig. S2G). This is in line with earlier reports defining asymmetric stem cell division as a feature of psoriatic skin (Charruyer et al., 2017, Li et al., 2020). Therefore, our data implies that during inflammation, *Fbln7* KO fast-cycling EpSCs undergo asymmetric division towards the epidermal suprabasal layers, while the number of total *Slc1a3-*tdT+basal cells remains unchanged.

### Epidermal stem cells transcriptome reveals genes modulated by fibulin 7 during inflammation

Bulk RNA-sequencing was performed on EpSCs isolated from the IFE of 1-year-old *Fbln7* WT and KO mice which had undergone 4 days of vas-control or IMQ treatment (Fig. 2, S3A). Principal component analysis depicts the transcriptome differences between WT vs KO EpSCs post-IMQ treatment (Fig. 2A), but not in vas-treated condition, signifying their differentially expressed genes (DEGs) in response to IMQ, which are illustrated in the volcano plot (Fig. 2B, table S1). Gene ontology (GO) analysis suggests enrichment of inflammatory pathways in KO such as LPS, TNF and IL-17 (Fig. 2C).

**Figure 2.**
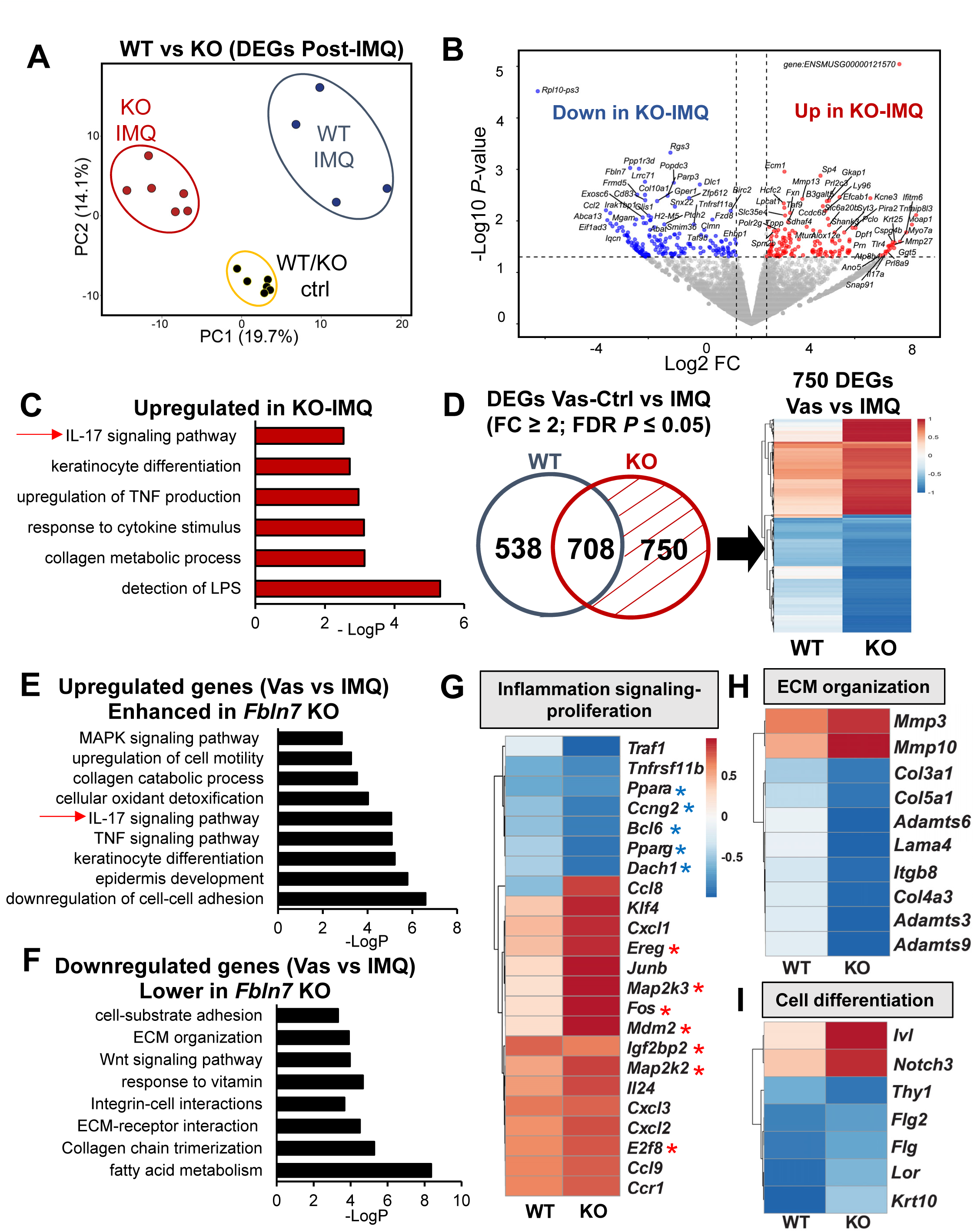
Transcriptomics of *Fbln7* WT vs KO dorsal skin epidermal stem cells in control and imiquimod-treated mice. Dorsal skin was treated with vas-control or IMQ as in figure 1A. FACS-sorted EpSCs (sorting strategy in figure S3A) were then subjected to bulk RNA-sequencing. n = 3 from WT or KO vas-control group, n = 4 from WT-IMQ group and n = 6 from KO-IMQ group. **(A)** Principle component analysis map constructed from the differentially expressed genes (DEG; fold change (FC) 1.5, *P* < 0.05)) between 2 groups, WT vs KO (post-IMQ) and **(B)** the corresponding volcano plot. FC, fold change. **(C)** Gene ontology (GO) from genes shown in the volcano plot as ‘Up in KO’ after IMQ application. **(D)** Transcriptome analysis from 4 groups (WT-vas vs WT-IMQ and KO-vas vs KO-IMQ) with FC ≥ 2 and FDR *P* ≤ 0.05. Venn diagram indicates 750 genes that are more significantly/highly regulated in the KO group upon IMQ treatment, which are then represented in the heat map (right panel). **(E)** Graphs describe GO of upregulated genes in the KO from the 750 gene pool and **(F)** the corresponding downregulated genes. Examples of these genes and their categories are represented in **(G, H, I).** All heat maps represent log2 FC from IMQ compared to control in WT or KO groups. Blue and red asterisks represent genes suppressing and promoting proliferation, respectively.

Subsequently, we performed a four-group DEG analysis that accounts for basal gene expression in vas-treated WT and KO compared to their respective IMQ-treated skin (Fig. 2D). A Venn diagram summarizes these DEGs and reveals that 750 genes were more strongly affected in KO, 708 genes were commonly modulated in both WT and KO, and 538 genes were preferentially regulated in WT (Fig. 2D, S3B, S3C; tables S2, S3, S4). The 750 KO-enriched genes included pathways related to keratinocyte differentiation and motility as well as IL-17 and TNF signaling that involves MAPK activation, whereas downregulated genes were associated with fatty acid metabolism, ECM organization, and Wnt signaling (Fig. 2E, F). Examples of these genes are shown in Fig. 2G – I, including inflammatory mediators linked to IL-17 signaling (such as *Ccl8*, *Cxcl1/2/3*, *Ereg*, *Fos*, *Junb*, *Mmp3/10* genes), proliferation-related genes, ECM remodeling components, and keratinocyte differentiation genes. Among the 708 commonly regulated genes, GO analysis showed similar pathway enrichment between WT and KO, with notable exceptions such as HIF-1 signaling pathway (Fig. S3D, S3E). Thus, this transcriptomic data indicates that both WT and KO EpSCs share broadly similar inflammatory responses to IMQ; however, loss of fibulin 7 may amplify this response, particularly through enhanced TNF and IL-17 signaling (Griffiths et al., 2021).

### Fibulin 7 mitigates IMQ- and IL-17A-mediated inflammatory signaling pathway

Our transcriptome analysis of post-IMQ *Fbln7* KO vs WT EpSCs revealed a subset of intensified inflammatory genes including *Fos* and *Junb* (Fig. 2D, G), which encode for AP-1 transcription factors downstream of MAPK signaling implicated in psoriasis pathogenesis (Guo et al., 2023, Hammouda et al., 2020). Consistent with prior reports showing elevated c-Jun N-terminal kinase (JNK) activity in human psoriatic epidermis (Takahashi et al., 2002), we assessed JNK phosphorylation in *Fbln7* WT and KO mouse epidermis after IMQ application. In the vas-control group, average phospho-JNK signal was low in basal cells of both *Fbln7* WT and KO (white arrowheads). Following IMQ treatment, WT epidermis exhibited moderate JNK activation, whereas KO epidermis showed a marked enhanced response (Fig. 3A, B). These findings suggest that fibulin 7 restrains JNK-driven inflammatory signaling in the epidermis.

**Figure 3.**
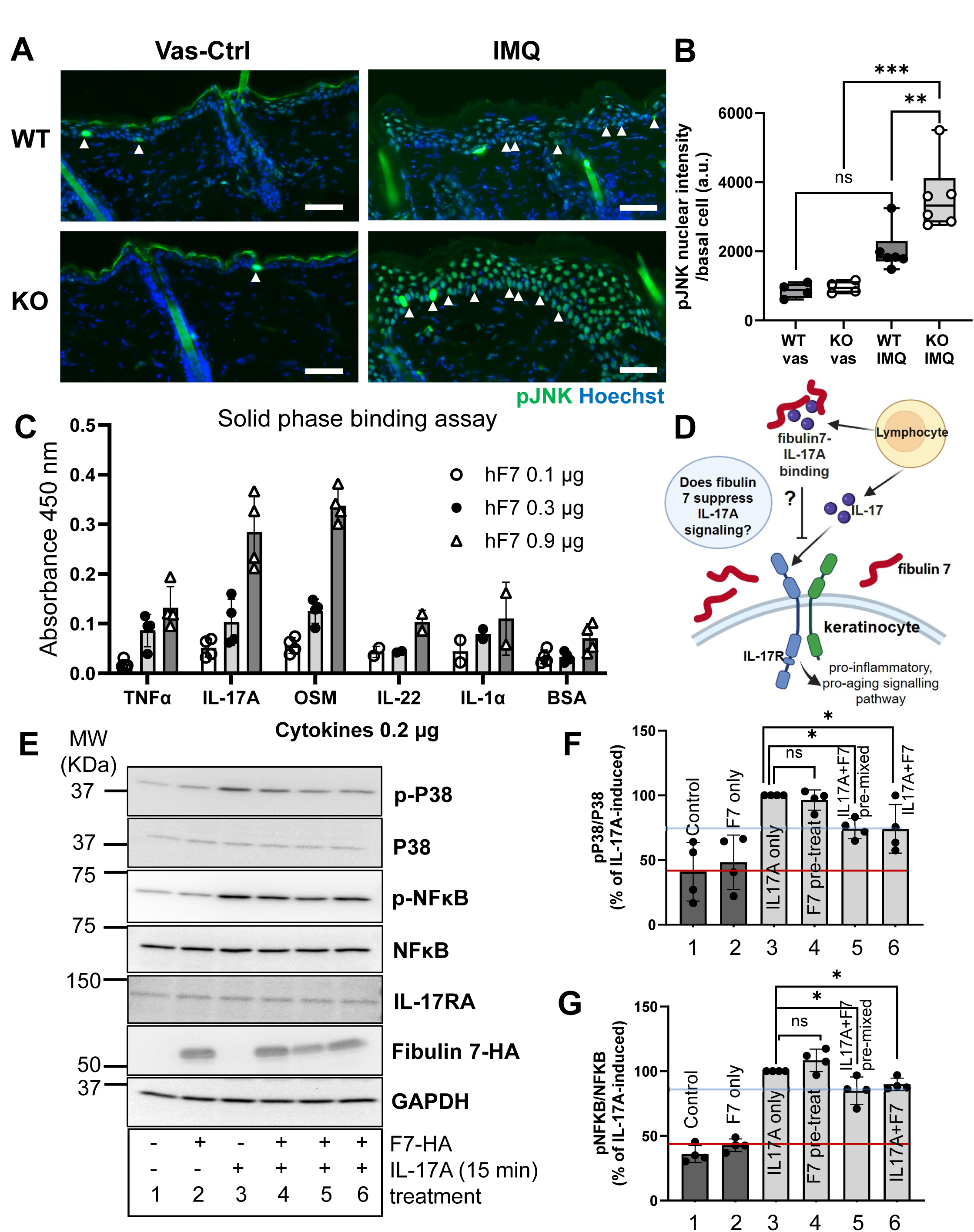
Fibulin 7’s interaction with IL-17A contributes to lower inflammatory signaling. **(A)** Immunofluorescence images of *Fbln7* WT and KO dorsal skin epidermis after 4 days of vas-control or IMQ application, showing nuclear phospho-JNK (p-JNK) and Hoechst nuclear counterstain. Examples of basal cells expressing p-JNK are pointed with white arrow heads. **(B)** Graph summarizes basal cells intensity for p-JNK (a.u., arbitrary unit). 2-way ANOVA, Tukey test. n = 4 for each vas-control group and n = 6 for each IMQ-treated group. **(C)** ELISA-based assay indicates the dose-dependent binding of soluble recombinant human fibulin 7 (hF7) to immobilized cytokines/bovine serum albumin (BSA) control at 200 ng/well. OSM, oncostatin M. Graph represents mean from 4 technical repeats, 2 independent experiments. **(D)** Hypothesis: illustration of fibulin 7 binding to IL-17A secreted by immune cells to reduce IL-17A interaction with its receptor IL-17R on the keratinocytes hence dampening its signaling. Drawing made in BioRender. **(E)** Immunoblots of primary human keratinocytes culture treated with (1) no treatment control, (2) HA-tagged fibulin 7 for 45 minutes, (3) IL-17A for 15 minutes, (4) fibulin 7 for 30 minutes before IL-17A addition for another 15 minutes, (5) fibulin 7 and IL-17A for 15 minutes (pre-mixed/binding for 30 minutes at 37^ο^C before adding to cells), and (6) fibulin 7 and IL-17A, simultaneously added to cells for 15 minutes. **(F, G)** Signals for phospho-p38 and phospho-NFκB were quantified from 4 independent experiments, normalized with their total protein levels and represented as a percentage of IL-17A-induced signal (set at 100%). Mann-Whitney tests. Bar graphs show mean ± S.D. * *P* < 0.05, ** *P* < 0.01, *** *P* <0.001, ns; not significant. Red and blue lines are respectively positioned on the approximate basal value on the graphs and reduction after adding fibulin 7.

To test whether fibulin 7 acts as a cytokine-binding ECM to modulate its signaling, we selected five candidate cytokines and tested their interaction with fibulin 7 (Fig. 3C, (Guilloteau et al., 2010, Rabeony et al., 2014)). Among them, fibulin 7 could bind to IL-17A and oncostatin M (OSM) in a dose-dependent manner (Fig. 3C). Although OSM exerts pro-inflammatory effects in human keratinocytes, its role in the IMQ-induced mouse model remains unclear (Pohin et al., 2016). In contrast, IL-17A is a central cytokine in both human and mouse psoriasis and IL-17A signaling in keratinocytes plays a major role in IMQ-induced psoriasis model (Moos et al., 2019), as well as inflammation in broader contexts (Huangfu et al., 2023, Sola et al., 2023). We therefore investigated whether fibulin 7 binding to IL-17A interferes with the IL-17A-induced activation of IL-17R and downstream signaling (Fig. 3D).

Primary human keratinocytes were briefly stimulated with IL-17A for 15 minutes to assess receptor-mediated activation of MAPK P38 and NF-κB in the presence or absence of fibulin 7 (Fig. 3E). Fibulin 7 alone did not affect signaling (Fig. 3E, lane 2), and pre-incubation of cells with fibulin 7 before IL-17A stimulation did not alter signaling (Fig. 3E, F, G; lane 4 vs lane 3). Fibulin 7 was reported to bind heparan sulfate proteoglycans on the cell surface (Sugiura et al., 2023, Tsunezumi et al., 2018) and our results here suggest that cell-associated fibulin 7 does not effectively sequester IL-17A. In contrast, when IL-17A was pre-mixed with fibulin 7 or when both were added simultaneously to cells, fibulin 7 significantly suppressed IL-17A-induced signaling (Fig. 3E, F, G; compare lane 3 vs lanes 5 and 6, Fig. S4A). It is plausible that fibulin 7 suppresses IL-17A signaling primarily by binding the cytokine prior to receptor engagement, rather than when immobilized at the cell surface.

Fibulin 7 also interacts with collagen IV, a major structural component of the BM (Lennon and Sherwood, 2025, Raja et al., 2022). Although the role of collagen IV in psoriasis remains unclear, activated T cells expressing α1β1 integrins can enter the epidermis via interactions with collagen IV in a human-mouse xenograft psoriasis model (Conrad et al., 2007). We then examined whether fibulin 7 modulates immune cell infiltration from the dermis into the epidermis in our mouse model (Fig. S4B). Flow cytometric analysis of pan-immune CD45+ cells indicated that, after 4 days of IMQ application, the number of epidermal immune cells was not significantly higher than vas-treated controls, and no significant difference was observed between KO-IMQ and WT-IMQ mice (Fig. S4C, S4D). Consistently, CD45 immunostaining confirmed immune cell recruitment primarily to the dermis and close to epidermal-dermal junction following IMQ administration, both in WT-IMQ and KO-IMQ mice (Fig. S4E). Subsequent characterization of CD3+ pan-T cells revealed sparse infiltration in the epidermis and dermis, indicating minimal T cell involvement during early IMQ-induced inflammation (Fig. S4F, (Tortola et al., 2012)).

Lastly, because STAT3 signaling can be indirectly activated by IL-17A (Furue et al., 2020) and plays a critical role in keratinocytes-driven IMQ-induced mouse model (Ravipati et al., 2022, Sano et al., 2005), we examined STAT3 activation. Immunostaining revealed that STAT3 is mainly expressed in the epidermal and hair follicle keratinocytes (Fig. S4F), and immunoblotting showed that phospho-STAT3 levels were increased in KO-IMQ mice, after normalization to total-STAT3 (Fig. S4G, S4H). Overall, these findings suggest that fibulin 7 modulates IL-17A-driven inflammatory signaling.

### Evaluation of cytokines/chemokines profiles in total skin tissue and cellular sources of fibulin 7 in psoriasis patients

To evaluate the broader cytokine landscape and its regulation by fibulin 7, we used whole-skin protein extracts from the IMQ-induced mouse model (Fig. 4A, B). Quantified data summarized in a heat map describes commonly upregulated inflammatory proteins (Fig. 4C, yellow highlight). Those that were more abundant in the KO skin compared to WT post-IMQ are shown in Fig. 4D and some candidates related to IL-17A signaling were further validated by western blotting. Quantification confirmed a significant increase in CXCL5 levels in IMQ-treated KO skin (Fig. 4E, F). IL-17A, IL-6 and CXCL2 showed similar upward trends, although these did not reach statistical significance (Fig. 4E, 4G – I).

**Figure 4.**
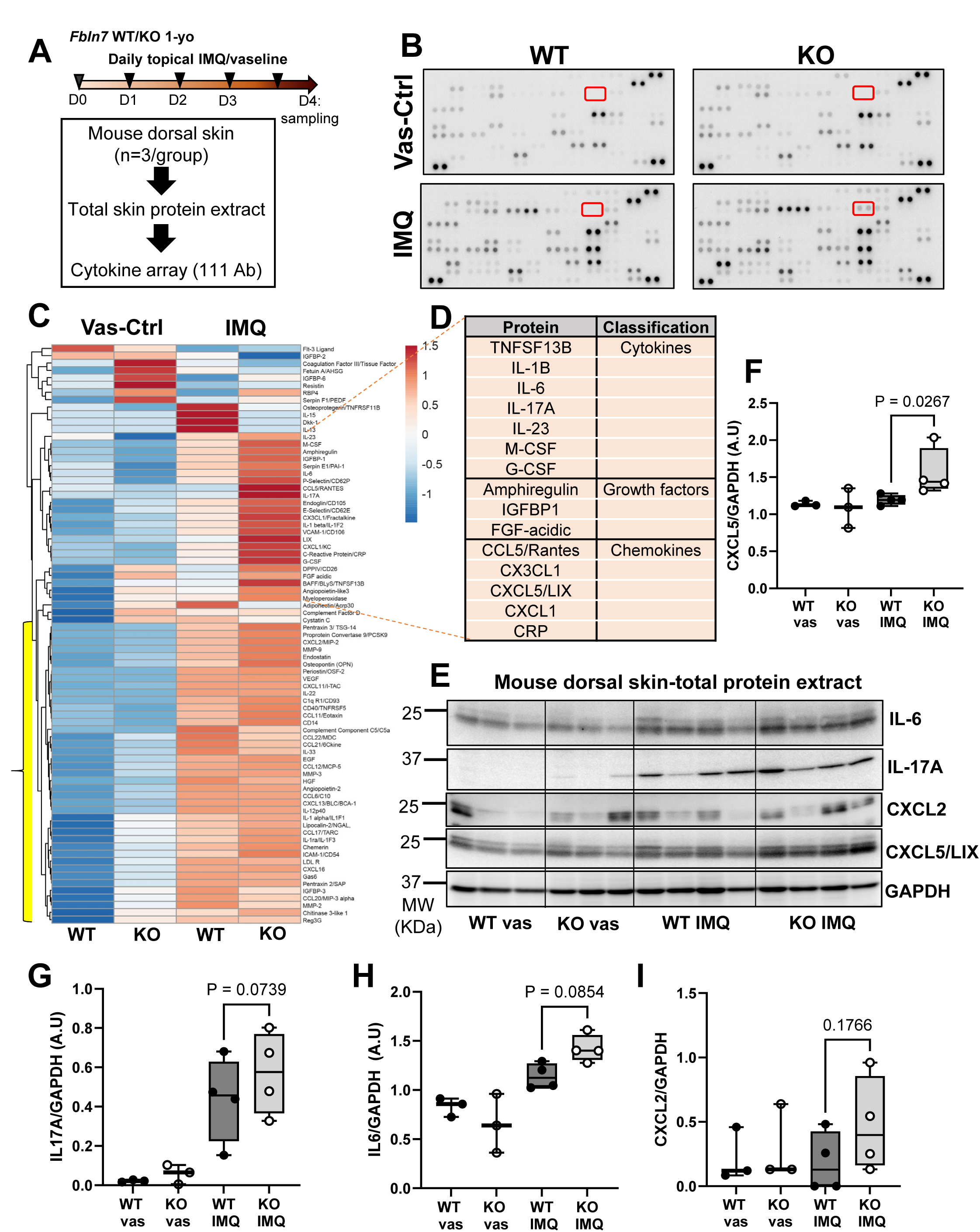
Inflammatory proteins abundance in *Fbln7* WT and KO total mouse skin extract. **(A)** The experimental setup for cytokine screening using whole mouse skin: timeline and number of mice used per group for protein extraction and binding to immobilized antibodies (Ab) array. **(B)** Chemiluminescent images of each Ab array membrane after binding with antigens from the total skin extract in each experimental group. Ab are presented as duplicate dots on the membrane and values were averaged for quantification presented in figure 5a. Red squares indicate signals from CXCL5/LIX antibody. **(C)** Heatmap summarizes changes in the level of cytokines and chemokines quantified from total skin extract of vas and IMQ-treated dorsal skin of *Fbln7* WT or KO mice subjected to a cytokine array, average of n = 3 per group. Scale bar, log2 protein abundance (arbitrary unit). **(D)** Table highlights some of the upregulated proteins in the KO-IMQ skin. **(E)** Immunoblots depicts protein abundance of IL-6, IL-17, CXCL2, CXCL5/LIX from an independent experiment. Each lane represents proteins from one mouse. **(F, G, H, I)** Quantification of cytokine/chemokine immunoblots normalized by GAPDH, from (e). 2-way ANOVA, fishers test.

To identify the primary cellular sources of fibulin 7 and its regulation in human psoriatic skin, we analyzed a public single-cell RNA-seq dataset (GSE173706). UMAP specifies the transcriptomes of 9 separate cell types (Fig. 5A) and dot plot confirms their marker gene expression (Fig. 5B). *FBLN7* was predominantly observed in melanocytes and fibroblasts (Fig. 5C). Interestingly, melanocytes are enriched in the basal layer of epidermal rete-ridges in human skin (Hirobe et al., 2021), consistent with the localization of fibulin 7 observed in Fig. S1D. In psoriatic lesions, melanocytes exhibited lower *FBLN7* expression compared to non-lesion skin, whereas fibroblasts expression was unaltered by disease state (Fig. 5D). Next, we compared *FBLN7* expression in total skin cells from paired psoriasis-lesion and non-lesion samples across 11 individuals. The resulting dot plot showed that 5 of 11 pairs displayed decreased *FBLN7* expression in lesional skin (Fig. 5E), supporting a potential anti-inflammatory role for fibulin 7.

**Figure 5.**
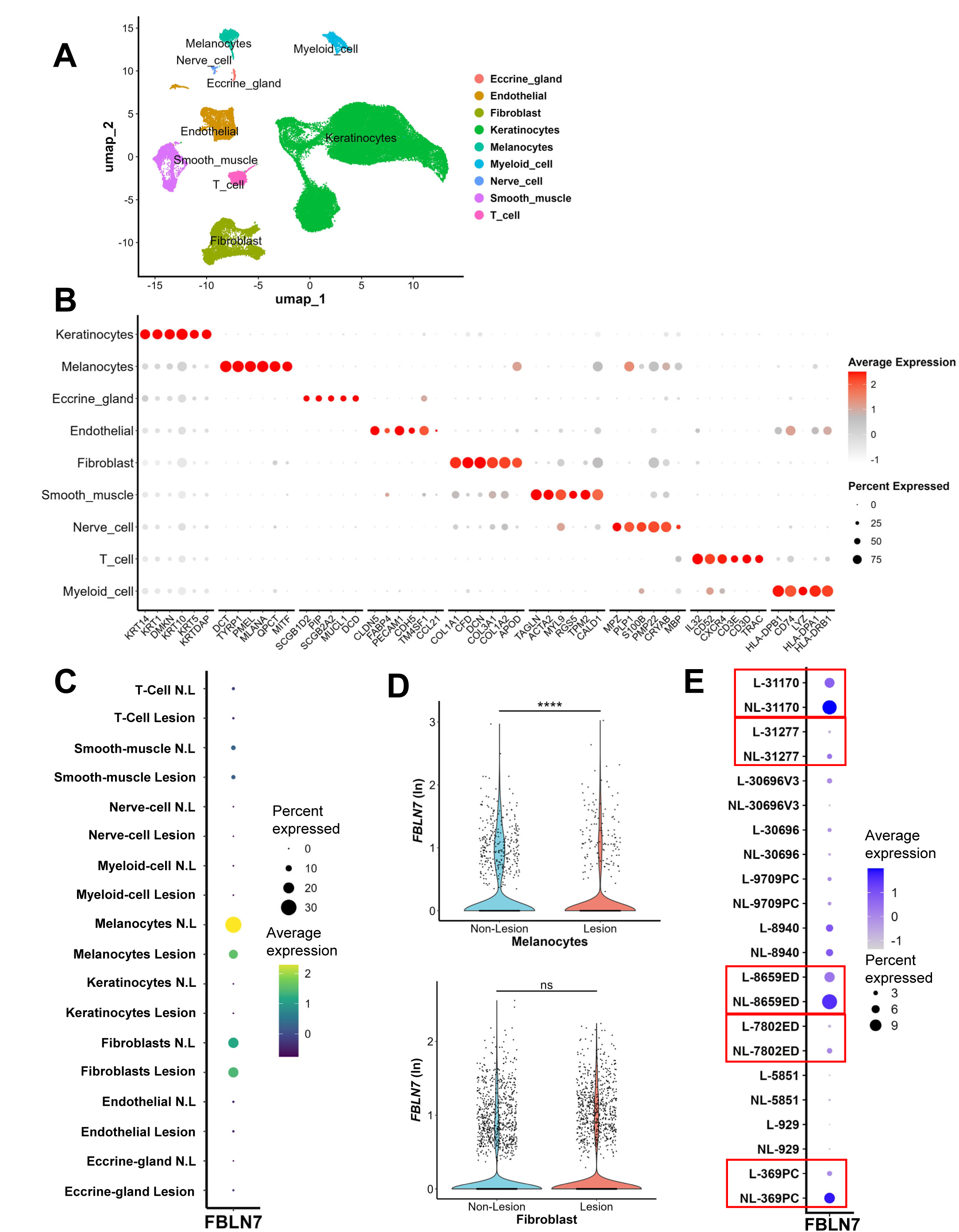
Cell types contribution to *FBLN7* expression in human psoriatic skin. **(A)** Human psoriasis scRNA-seq public dataset GSE173706 (n =11 pairs of non-lesion and lesions) was used to generate UMAP plot containing 59593 cells separated into 9 cell-types. **(B)** Dot plot describes the 9-cell type’s distinction based on their gene markers. **(C)** *FBLN7* expression per cell type, N.L; non-lesion. **(D)** *FBLN7* plots from melanocytes and fibroblasts. ns, not significant. **** adj*. P* < 0.0001. **(E)** Analysis of *FBLN7* expression in all cells of 11 paired patient skin data. L; psoriasis lesion, NL; non-lesion. Red boxes indicate pairs with lesion cells expressing lower *FBLN7* than non-lesion cells.

### *FBLN7* expression negatively correlates with human psoriasis lesions and their associated genes

To further evaluate the involvement of fibulin 7 in human psoriasis cohorts, we analyzed 3 independent transcriptomic datasets comparing lesion and non-lesion skin (Fig. 6, S5), encompassing a total of 176 lesion, 171 non-lesion, and 85 normal skin samples. In all 3 datasets, *FBLN7* mRNA levels were lowest in lesional skin compared to non-lesion or normal skin (Fig. 6A, S5A), consistent with our findings in Fig. 5E. Analysis of common DEGs (984 upregulated and 630 downregulated in psoriasis lesions) revealed that *FBLN7* expression was broadly and inversely correlated with these gene sets (Fig. 6B, S5B, S5C; Table S5). In GSE13355, about 50% of common psoriasis-DEGs showed significant inverse correlations with *FBLN7* (Fig. 6B), while 30-45% of genes in GSE30999 showed similar associations (Fig. S5C). Representative genes and their correlation coefficients to *FBLN7* are shown in Fig. 6C, highlighting classical enrichment in pathways related to inflammatory signaling, DNA repair, keratinocyte differentiation, protein synthesis, protease-mediated degradation, ECM remodeling, and growth factor signaling. Moreover, scatter plot analysis demonstrated strong inverse relationship between *FBLN7* expression and key psoriasis-related genes across lesion and non-lesion samples (Fig. S5D, S5E).

**Figure 6.**
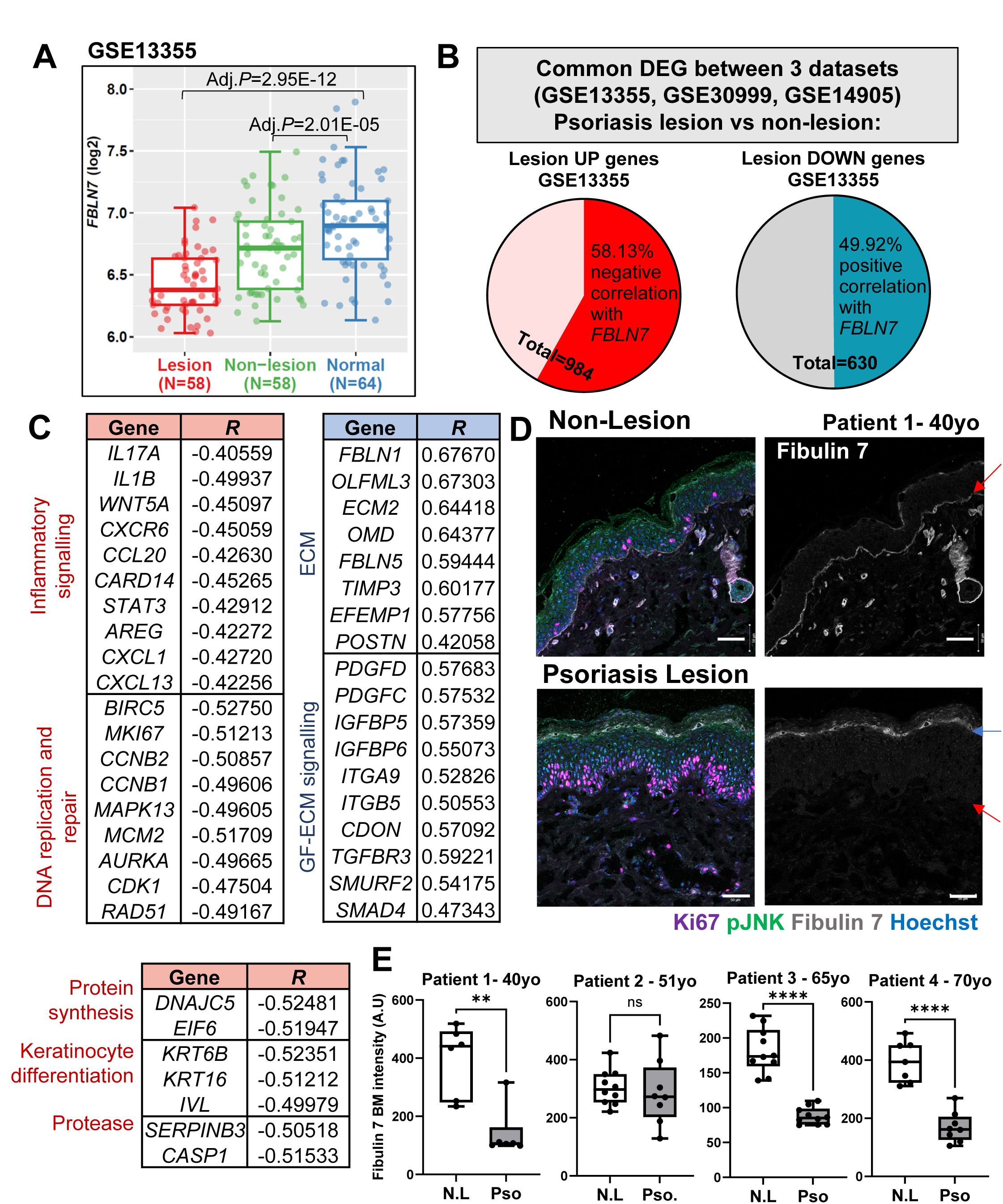
*FBLN7* expression is lower in human psoriatic skin lesion and is negatively correlated with psoriasis-associated genes in larger cohorts. **(A)** *FBLN7* mRNA level from public dataset GSE13355. **(B)** Venn diagrams illustrate the proportion of commonly regulated psoriasis-related genes between 3 datasets (1.5-fold change and *adj. P* < 0.05; lesion vs non-lesion) that have significant correlation with *FBLN7* expression (Pearson correlation *R* cut-off 0.4 or -0.4, adj *P*-value < 0.05). **(C)** Gene examples upregulated in psoriasis lesion with negative correlation to *FBLN7* (marked in red) and those downregulated in lesion with positive correlation to *FBLN7* (marked in blue). GF, growth factor. **(D)** Immunostaining images from a 40-year-old patient skin showing fibulin 7 at the basement membrane (right panel, red arrows) which decreased in the lesion areas. Scale bars, 50 μm. **(E)** Quantification of fibulin 7 abundance at the BM from (d) and 3 other patients aged 51, 65 and 70-years-old. Each dot represents an average value from one image field. N.L, non-lesion; Pso, psoriasis lesion. Unpaired Welch T-tests. ** *P* < 0.01, **** *P* < 0.0001, ns; not significant.

Finally, we examined fibulin 7, Ki67 and phospho-JNK in skin samples from 4 Japanese patients with psoriasis (aged 40 – 70 years old). In contrast to young normal skin, which showed fibulin 7 enrichment at the rete-ridges (Fig. S1D), non-lesion skin from older individuals exhibited a flatter epidermis with a fine uniform distribution of fibulin 7 along the BM (Fig. 6D, S6A – S6C). Notably, BM fibulin 7 levels were reduced in the lesion epidermis of 3 patients compared to the paired non-lesion (Fig. 6D, 6E, S6A – S6C). These changes were accompanied by increased Ki67+ cells and elevated phospho-JNK, indicating enhanced keratinocyte proliferation and inflammatory signaling in lesion epidermis.

## Discussion

Various tissue stem cells are equipped with intrinsic defense mechanisms such as expression of interferon-stimulated genes at steady state for anti-pathogen alertness (Poirier, 2025). However, under inflammation and cytokine surges during injury, infection, or disease, the autonomous stem cell function and tissue repair can be compromised. Identification of additional defense mechanisms within the stem cell niche that guard them during inflammation is necessary.

The ECM in the vicinity of stem cells could play such a role (Donnelly et al., 2024), despite extensive remodeling during intense inflammation. In psoriasis, structural BM modifications in the epidermis have been consistently reported, such as frequent gaps and irregular thickening accompanied by augmented collagen IV, laminins, matrix metalloproteases (MMPs) in both human and IMQ-induced mouse skin (Dong et al., 2024, McFadden and Kimber, 2016, Natsumi et al., 2018, Noddeland et al., 2024). Likewise, collagen I and lysyl oxidase are over-expressed along the dermal-epidermal junction, increasing overall stiffness, mechanotransduction, and basal cell proliferation (Balsini et al., 2025, Jiang et al., 2024). Unlike structural ECM, matricellular proteins such as fibulins dynamically move within the BM structural scaffold to prevent tissue damage during mechanical changes (Keeley et al., 2020). It is tempting to speculate that fibulin delivery supports BM integrity and reduces stiffness, thereby easing epidermal hyperproliferation in a psoriasis setting.

Our findings showcase fibulin 7 as an active participant in the BM to attenuate inflammatory response by binding to cytokines such as IL-17A and OSM, thus limiting EpSCs exposure to excessive inflammatory signals during psoriasis development and possibly in aging skin, where IL-17A is also aberrantly expressed (Li et al., 2019, Raja et al., 2022, Sola et al., 2023). As a result, in the absence of *Fbln7* during IMQ or cytokine induction, IL-17 signaling mediators such as JNK and p38 MAPKs were more activated in keratinocytes and EpSCs, enhancing relevant IL-17 target genes. Interestingly, AP-1 transcription factors were among the higher IMQ-promoted genes in *Fbln7* KO EpSCs, raising a question of whether these KO cells would be more sensitized to future inflammatory insults (Larsen et al., 2021).

Of note, fibulin 7 appeared to suppress IL-17A – p38 pathway better than IL-17A – NF-κB pathway, implying that its effect may not be solely mediated by cytokine sequestration. IL-17A homodimers bind to their receptor complex IL-17RA and IL-17RC (Huangfu et al., 2023). However, it also binds to IL-17RD, which is necessary for activation of p38/JNK but dispensable for NF-κB pathway (Mellett et al., 2012, Su et al., 2019). It is probable that IL-17A – fibulin 7 complex primarily hinders IL-17A interaction with IL-17RD, therefore weakening p38 signaling more than NF-κB, as indicated by p-p38 vs p-NF-κB in primary keratinocytes and IL-17A target genes in EpSCs of *Fbln7* KO-IMQ that overlap with downregulated genes in IL-17RD KO keratinocytes (Su et al., 2019)). NF-κB activity is central in inflammatory pathways, but p38 signaling in keratinocytes is also substantial in mediating IL-17 or IMQ-associated inflammation (Zheng et al., 2024).

In addition to signaling and transcriptomic observations, cytokine screening suggests that the chemokine CXCL5 is elevated in *Fbln7* KO skin treated with IMQ. CXCL5 is expressed by damaged or stressed epithelial cells to recruit neutrophils (Mathur et al., 2019, Moreau et al., 2021). CXCL5-neutrophil axis was reported to impair hair follicle stem cells (HFSCs) differentiation towards epidermal lineage during injury repair and worsen psoriatic inflammation in diabetic obese mice (Mathur et al., 2019, Shimoura et al., 2018). Neutrophils infiltrate the epidermis early during psoriasis development (Flutter and Nestle, 2013, Katayama, 2018) and were shown to be dependent on IL-17RA signaling in keratinocytes (Moos et al., 2019). It is possible that through CXCL5 neutrophils recruitment is enhanced upon IMQ induction in *Fbln7* KO skin, which could be a topic of future investigation.

In conclusion, our study identifies fibulin 7 as a protective ECM component that functions through multiple mechanisms. Firstly, fibulin 7 maintains fast-cycling EpSC population in middle-aged skin. Although the role of EpSC population balance in disease initiation or progression is unknown, a recent report suggests that during acute inflammation-induced epidermal hyperplasia, fast-cycling EpSCs and its differentiation was shifted towards the slow-cycling lineage by Wnt signaling suppression (Phung et al., 2026). Our *Fbln7* KO EpSCs transcriptome similarly indicated lower Wnt signaling post-IMQ and fewer *Slc1a3*+ fast-cycling EpSCs were detected by tdTomato-labeling in the KO epidermis, in line with our previous report as well (Raja et al., 2022).

Secondly, fibulin 7 regulates IL-17A signaling by direct binding to the ligand and attenuating downstream signaling. Moreover, we demonstrated an inverse relationship between *FBLN7* expression and psoriasis lesions in human cohorts. We propose that fibulin 7 acts as a barrier component in BM, restricting IL-17A access to EpSCs and limiting inflammatory responses in IMQ-induced epidermal hyperplasia. Given the central role of IL-17A signaling in a wide range of pathological conditions (Huangfu et al., 2023), these findings warrant further investigations.

### Study limitations and future directions

In this study fast-cycling EpSCs were labeled utilizing the *Slc1a3*-CreER-tdTomato system. However, Slc1a3 is expressed not only in EpSCs within the IFE but also in HFSC, leading to concurrent labeling of both populations (Reichenbach et al., 2018, Sada et al., 2017). HFSCs are also activated by IMQ (Amberg et al., 2016) and during inflammation, HFSCs can migrate to the IFE and adopt EpSCs-like properties to facilitate tissue repair (Ge et al., 2017). Although CXCL5 in the *Fbln7* KO may limit such lineage conversion following inflammation (Mathur et al., 2019), it remains likely that a subset of *Slc1a3*-tdT+ cells in the IFE originates from the hair follicle after IMQ-induced inflammation. Future studies could address the molecular domain(s) of fibulin 7 responsible for cytokine binding and to determine whether its inhibitory effect can be enhanced by modulating interactions with cell surface proteins or by engineering smaller-sized fibulin 7 variants with improved cytokine sequestration capacity to facilitate therapeutic application.

## Materials and methods

### Mice

Experiments were done following the guidelines of Animal Experiment Committee at the University of Tsukuba under standard housing conditions and 12-hour dark/light cycles. *Fbln7* KO mice (129SvEv; C57BL/6J) was previously established (Tsunezumi et al., 2018). Slc1a3^CreER^ and Rosa-tdTomato reporter mice (both C57BL/6J; The Jackson Laboratory, no 012586 and 007905, respectively) were crossed and the resulting mice were introduced into the *Fbln7* KO mice for lineage tracing. Experiments include both female and male mice.

### Imiquimod and tamoxifen administration

62.5 mg of 5% IMQ cream (Perrigo) or control Vaseline was topically applied daily over 4 days to 1-year-old mice (12-13 months old) dorsal skin after hair removal by shaving and depilation cream. Mice were euthanized on the fifth day and skin samples collected. For lineage tracing, Tamoxifen (Sigma) was delivered intraperitoneally to 1-year-old mice daily at 100 μg/g body weight for 5 days. After 3 weeks daily topical IMQ or vas treatment was started as described above.

### Human skin samples

Normal human abdominal skin from 25 to 35-years-old individuals (SK0548, SK0163, SK0558, SK0545) were supplied by CTI-Biotech (Lyon, France). Human psoriasis skin (lesion and non-lesion) samples were obtained from Hokkaido University. All human tissue was fresh frozen in OCT blocks and were thin-sectioned to 10-μm slices prior to immunostaining.

### Ethics statement

Animal experimentation protocol has been approved by the Institutional Animal Experiment Committee (University of Tsukuba, no 25-318). Normal human skin samples from CTI-Biotech (Lyon, France) and human primary keratinocytes from BIOPREDIC International (Saint-Grégoire, France) were collected with written, informed consent from donors in accordance with the European ethical standards. The study using psoriasis patients’ lesion and non-lesion skin was approved by the ethical committees from both University of Tsukuba (R05-168) and Hokkaido University (14-063). The study participants have given their written informed consent.

### Data availability statement

Data is available within the manuscript and references to transcriptomic data deposition are available in the materials and method.

## Acknowledgements

We thank the Animal Resource Center (University of Tsukuba) for providing high-quality animal care, M. Ho and T. Zama for administrative help, M.Sanaki-Matsumiya for technical advice, E. Motoyama, A.A. Sheikh, M. Tanabe and T.K.N. Nguyen for technical help. This work was supported by Grant-in-aid-for Scientific Research (C) (JP23K07737 to E.R), Grant-in-Aid for Research Activity Start-up (JP20K22659 to E.R), Lydia Òleary Memorial Pias Dermatological Foundation/Elastin molecular research grant 2022, the Cooperative Research Project Program of Life Science Center for Survival Dynamics, Tsukuba Advanced Research Alliance (TARA Center) (to A.S and J.T) and JST SPRING (to Narenmandula).

## Authors contributions

Conceptualization: ER, HY; Formal analysis: ER, TM, N, KE; Funding acquisition: ER, AS, JT; Investigation: ER, TM, ASM-SH, FW; Methodology: ER, AS, KA, KK; Project administration: ER, HY; Resources: JT, YW, KN, AS; Supervision: ER, HY; Validation: ER, TM; Visualization: ER, N, KE; Writing - original draft: ER; Writing – review and editing: ER, HY, AS, KN, KA, JT, KE.

## Declaration of generative AI use

ChatGPT was used to assist in language editing (to enhance clarity and conciseness). The authors carefully reviewed and take full responsibility for the edited version of the manuscript.

## Abbreviations

IMQ: imiquimod
vas: Vaseline
ECM: extracellular matrix
EpSCs: epidermal stem cells
IFE: interfollicular epidermis
BM: basement membrane
WT: wild type
KO: knock-out
IL-17: interleukin-17.

