## Supplementary Figures and Methods for "The epidermal stem cell-supporting matricellular protein fibulin 7 modulates skin inflammatory response in a psoriasis model"

[1]Life Science Center for Survival Dynamics, Tsukuba Advanced Research Alliance (TARA), University of Tsukuba, Japan, [2]Graduate School of Comprehensive Human Sciences, University of Tsukuba, Japan, [3]Leibniz Research Center for Working Environment and Human Factors, Dortmund University, Germany, [4]International Research Center for Medical Sciences (IRCMS), Kumamoto University, Kumamoto, Japan, [5]Molecular Pathology Division, Kanagawa Cancer Center Research Institute, Kanagawa, Japan, [6]Institute of Medicine, University of Tsukuba, Japan, [7]Department of Dermatology, Faculty of Medicine and Graduate School of Medicine, Hokkaido University, Japan, [8]Division of Skin Regeneration and Aging, Medical Institute of Bioregulation, Kyushu University, Fukuoka, Japan, [9]Department of Pharmacy, Varendra University, Bangladesh, [10] Faculty of Medicine, University of Tsukuba, Tsukuba, Japan.

*Corresponding authors:

Erna Raja, Ph.D.

Hiromi Yanagisawa, M.D., Ph.D.

**This file includes the following:**

Supplementary Materials and Methods

Supplementary Figures S1 - S6

**Hematoxylin and eosin staining**

H&E staining was done on 10-μm dorsal skin slices from fresh frozen tissue in OCT blocks as previously described (Raja et al., 2022).

**Tissue section immunostaining**

Immunostaining procedures for both mouse dorsal skin or human skin sections were as described in (Dumrongphuttidecha et al., 2025). Primary antibodies were: rabbit anti-K14 (1:1000, BioLegend, 905304), HiLyte-fluor-555-conjugated (Dojindo) monoclonal mouse anti-fibulin 7 (1:800, (Raja et al., 2022)), rabbit anti-laminin (1:200, abcam, ab11575), guinea pig anti-Slc1a3 (1:100, Frontier Institute, GLAST-GP-Af1000), rat anti-Ki67 (1:200, eBioscience, 14-5698), rat anti-CD45 (1:400, BD Biosciences, 550539), rat anti CD-3 (1:200, BD Biosciences, 555273), rabbit anti-phospho JNK (1:200, Abcam, Ab124956) and rabbit anti-STAT3 (1:200, Cell Signaling, 30835). Secondary antibodies (Alexa 488, 546, 647, Invitrogen) were used at 1:200 dilution together with Hoechst nuclear stain (Sigma, B2261) before mounting.

**Whole-mount immunostaining**

Tissue was processed according to protocol in (Wong et al., 2021) with some adaptations. Dorsal skin pieces (about 5 $mm^2$ each) were incubated in primary (rabbit anti-K14, 1:500) and secondary (Alexa goat anti-rabbit 488, 1:200) antibodies for 2 days each at 4°C with shaking, and washing was done after each antibody incubation. Then Hoechst nuclear staining (1/1000) was performed overnight at 4°C followed by clearing. Tissue was first transferred to 30% glycerol/PBS solution for 1 hour at room temperature (RT) then into

*RapiClear* 1.52 solution (Sunjin Lab) for 5 to 7 hours at RT and mounted on a concave slide with the same clearing solution.

**Imaging and quantification**

Images were observed and captured by Zeiss Axio Imager Z2 microscope and Zeiss LSM980 confocal microscope with corresponding Zen software. Adjustments were made using Adobe photoshop 2018 or ImageJ software (NIH) prior to quantifications. ImageJ was employed to quantify skin thickness, phospho-JNK signals and Ki67+ cells with the following details. Average epidermal or dermal thickness were calculated from skin area/skin length (https://hookelabs.com/services/cro/psoriasis/imiquimod/) in 3-5 images per mouse. For fibulin 7 BM intensity measurement, the BM area was demarcated and intensity in the region of interest (ROI) was measured as average values per area. Similarly, phospho-JNK intensity was quantified from 100-200 basal cell nuclei/mouse. Ki67+ cells were scored from about 100-1000 tdTomato+ cells in 8-18 images per mouse. Additionally, tdTomato+ basal cells were analyzed and counted from wholemount immunostaining images using Imaris 10 software (Oxford Instruments).

**Primary keratinocytes culture**

Human adult (27-years-old) primary keratinocytes (KER110004, BIOPREDIC International) were seeded with laminin coating (iMatrix-511 silk, Nippi; pre-mixed 3 μL/ml of cell suspension) and cultured in CnT-PRIME medium (CELLnTEC). Cells detachment were induced by 1:2 ratio of Accumax to Accutase (both from Innovative Cell Technologies) for 10-15 minutes at RT. For experiments, cells up to passage 6 were utilized and seeded on 12-well-plates with a density of 50,000 cells/well. They were maintained in growth medium for 1-2 days before changing into experimental homeostasis medium CnT-PR-H (CELLnTEC) for 1 day. Next, 1.5 μg of recombinant human fibulin 7 (8530-FB, R&D Systems) and 250 ng/mL of recombinant human IL-17A (200-17, PeproTech) were added to the cells with combinations as indicated in figure legend.

**Solid phase binding assay**

This ELISA-based assay was performed as previously described in (Raja et al., 2022). 200 ng of cytokines (TNFα, IL-17A, Oncostatin M, IL-22, IL-1α, all from PeproTech) or BSA control (Sigma) control in bicarbonate buffer were added on 96-well-plate overnight at 4°C (as solid phase). The next day binding of 0.1 to 0.9 μg of HA-tagged human recombinant fibulin 7 (8530-FB, R&D Systems) was tested.

**Cytokine screening**

Proteome profiler kit (Mouse XL Cytokine Array Kit, R&D Systems) was utilized according to manufacturer's recommendations. Mouse dorsal skin was collected from 3 mice/group, snap-frozen in liquid $N_2$ and pulverized with mortar/pestle before adding lysis buffer (100 μL/10 mg tissue, 1% Triton-X-100 in PBS supplemented with protease inhibitor cocktail (Thermo Fisher Scientific). 180 μg of total protein lysate/group was incubated with the immobilized antibodies on a nitrocellulose membrane.

**Western blotting**

Mouse dorsal skin was pulverized and lysed in lysis buffer 17 (R&D Systems) containing protease and phosphatase inhibitor cocktails (Wako and Nacalai Tesque, respectively) at 150 μL/10 mg tissue. Keratinocytes were lysed in TNE buffer (1% IGEPAL-CA 630, 1 mM EDTA pH 8.0, 150 mM NaCl, 20 mM Tris pH 8.0). Protein concentration was measured by Pierce detergent compatible Bradford reagent and absorbance read with microplate reader (Varioskan LUX, Thermo Fisher Scientific). Equal amount of proteins was loaded into SDS-PAGE gels (Bio-Rad) and blotted onto PVDF membranes (Millipore). Membranes were incubated with primary antibodies in 0.1% TBS-Tween20 at 4°C overnight and secondary antibodies for 1 hour at RT prior to chemiluminescence detection (ATTO). Membrane washing was performed at 3x10 minutes after each antibody reaction. List of primary antibodies (diluted at 1/1000-2000) is as follows; anti-CXCL5/LIX (MAB433), anti-CXCL2/MIP2 (MAB452), anti-IL-6 (MAB406-SP), all from R&D Systems; anti-GAPDH (2118S), anti-IL-17A (13838S), anti-HA-tag (3724S), anti-phospho-Ser536-NFκB (3033S), anti-STAT3 (30835), anti-phospho-Tyr705-STAT3 (9145), anti-phospho-Thr180/Tyr182-p38 (9211S), anti-p38 (9212S), all from Cell Signaling; anti-NFκB (16502), anti-phospho-Thr183/221-JNK1/2/3 (Ab124956), both from Abcam; anti-IL-17RA (sc376374, SantaCruz).

**FACS isolation and analysis**

Mouse dorsal skin post-IMQ or vas treatment were excised and processed as before (Raja et al., 2022). For EpSCs isolation, epidermal cell suspension was stained with the following antibodies and dilutions for 30 minutes on ice: anti-Sca1-FITC (1:100, 557405) and anti-α6-integrin-PE (1:150, 555736), both from BD Pharmingen; anti-CD34-biotin (1:50, eBioscience, 13-0341-85); anti-CD31-biotin (102404), anti-CD117-biotin (105804), anti-CD45-biotin (103104), and anti-CD140a-biotin, all diluted at 1:200, from BioLegend; lastly, Streptavidin-APC (1:100, BD Biosciences, 554067). 7-AAD (1:300, 559925, BD Pharmingen) was added to stain and exclude dead cells. For epidermal CD45+ cell analysis, cell suspension was stained with anti-CD45-biotin, Streptavidin-

APC and 7-AAD. Cells were analyzed and sorted using FACS Aria flow cytometer and FlowJo software (both from BD Biosciences).

**Mouse cells RNA-sequencing and analysis**

Freshly sorted 150,000 EpSCs were directly suspended in Trizol (Ambion) and submitted to Azenta for RNA-quality control check and sequencing (Project 60-1104570568). FASTAQ raw data conversion and gene expression analysis were conducted in CLC Genomics 22 software (Qiagen). Online tools such as Venny 2.1 (https://bioinfogp.cnb.csic.es/tools/venny/), ClustVis (https://biit.cs.ut.ee/clustvis/) (Metsalu and Vilo, 2015) and Metascape (https://metascape.org/) (Zhou et al., 2019) were used for construction of Venn diagrams, PCA-heatmaps and GO analysis, respectively. Volcano plot was made using R (ver. 4.3.2) and ggplot2 (ver. 4.0.2). Sequencing data is stored at Gene Expression Omnibus (GEO) GSE319787.

**Human psoriasis public datasets analysis**

Publicly available Affymetrix Human Genome U133 Plus 2.0 Array data were downloaded from GEO data repository, including raw CEL files and metadata. GSE13355 dataset included data from 58 psoriatic patients (one punch biopsy from lesional skin and one from non-lesional skin) and 64 normal healthy controls (Nair et al., 2009). GSE30999 included data from skin biopsies collected from 85 patients with moderate-to-severe psoriasis (lesion and matched non-lesional skin) (Suarez-Farinas et al., 2012). GSE14905 included data from skin biopsy samples from 21 healthy donors, 33 psoriasis patients (28 matching lesions and non-lesions; 5 only lesions) (Yao et al., 2008). Statistical programming language R (ver. 4.2.3) was used for statistical analyses and graphic visualizations. The raw CEL files were normalized with Robust multiarray analysis (RMA) (Irizarry et al., 2003). Differentially expressed probesets were identified for each dataset using limma (Linear Models for Microarray Data) (cutoff: 1.5-fold change and false discovery rate (FDR) adjusted $P < 0.05$) (Smyth, 2004), and the overlap between the three datasets was next determined, resulting in 2328 common differentially expressed probesets, corresponding to 1614 differentially expressed genes (DEGs) (excluding *FBLN7*) after removal of non-annotated genes. In case of multiple probe sets per gene, the ones satisfying the cutoff criteria were retained. Data visualization was made in R package ggplot2 (ver.3.5.2). Correlation coefficients were calculated in the R function cor.test.

Public scRNA-seq dataset (GSE173706) (Ma et al., 2023) containing 11 pairs of psoriatic skin and matching peripheral normal skin (PP/PN-30696, 30696V3, 31170, 31277, 369PC, 5851, 7802ED, 8659ED, 8940, 929, 9709PC) were re-analyzed. Expression matrices were analyzed in R (ver.4.4.2) using Seurat package (ver. 5.2.1). After Seurat object was created quality control was conducted to select single cells with ≤ 20% mitochondrial genes expression, gene expression between 200 and 5000 genes (Stuart et al., 2019). Following normalization, data scaling and Principal Component Analysis (PCA) were conducted according to standard Seurat workflows. To correct for batch effects, datasets were integrated using the reciprocal PCA (RPCA) method.

Following integration, cells were embedded into a two-dimensional space using Uniform Manifold Approximation and Projection (UMAP) based on the first 30 principal components (PCs). To further refine data quality, we removed low-quality clusters, defined as those with a median nFeature_RNA < 800 or a median percent.mt > 8%. The remaining 59,593 high-quality cells were subsequently re-clustered. Cell type annotation was performed manually using established marker genes, including: Keratinocytes (*KRT14, KRT1, DMKN, KRT10, KRT5, KRTDAP),* Melanocytes (*DCT, TYRP1, PMEL, MLANA, QPCT, MITF),* Fibroblasts (*COL1A1, CFD, DCN, COL3A1, COL1A2, APOD),* Endothelial cells (*CLDN5, FABP4, PECAM1, CDH5, TM4SF1, CCL21),* Smooth muscle cells (*TAGLN, ACTA2, MYL9, RGS5, TPM2, CALD1),* T cells (*IL32, CD52, CXCR4, CD3E*), Myeloid cells (*HLA-DPA, HLA-DPB1, CD74, LYZ*), Eccrine glands (*SCGB1D2, PIP*), and Nerve cells (*MPZ, PLP1, S100B*). Visualization of cell clusters and marker gene expression was performed using ggplot2 (v3.5.2) and patchwork (v1.3.2). Differential gene expression analysis between lesional and non-lesional samples was conducted using the default Wilcoxon rank-sum test via the FindMarkers function implemented in the Seurat package.

**Statistics**

All statistical analyses were conducted in GraphPad Prism 10, except transcriptomic data. Shapiro-Wilk test was performed to analyze data normality and when normality could not be assumed, non-parametric test was performed. Statistical tests were employed to analyze differences between 2 groups only (unpaired T-tests), multiple groups and multiple comparisons (two-way ANOVA then Tukey test), or multiple groups but pairwise comparison only (two-way ANOVA then Fishers test).

Dumrongphuttidecha T, Ishikawa M, Cabezas-Wallscheid N, Sada A. Retinoic Acid Signaling Alters the Balance of Epidermal Stem Cell Populations in the Skin. J Invest Dermatol 2025.

Irizarry RA, Hobbs B, Collin F, Beazer-Barclay YD, Antonellis KJ, Scherf U, et al. Exploration, normalization, and summaries of high density oligonucleotide array probe level data. Biostatistics 2003;4(2):249-64.

Ma F, Plazyo O, Billi AC, Tsoi LC, Xing X, Wasikowski R, et al. Single cell and spatial sequencing define processes by which keratinocytes and fibroblasts amplify inflammatory responses in psoriasis. Nat Commun 2023;14(1):3455.

Metsalu T, Vilo J. ClustVis: a web tool for visualizing clustering of multivariate data using Principal Component Analysis and heatmap. Nucleic Acids Res 2015;43(W1):W566-70.

Nair RP, Duffin KC, Helms C, Ding J, Stuart PE, Goldgar D, et al. Genome-wide scan reveals association of psoriasis with IL-23 and NF-kappaB pathways. Nat Genet 2009;41(2):199-204.

Raja E, Changarathil G, Oinam L, Tsunezumi J, Ngo YX, Ishii R, et al. The extracellular matrix fibulin 7 maintains epidermal stem cell heterogeneity during skin aging. EMBO Rep 2022;23(12):e55478.

Smyth GK. Linear models and empirical bayes methods for assessing differential expression in microarray experiments. Stat Appl Genet Mol Biol 2004;3:Article3.

Stuart T, Butler A, Hoffman P, Hafemeister C, Papalexi E, Mauck WM, 3rd, et al. Comprehensive Integration of Single-Cell Data. Cell 2019;177(7):1888-902 e21.

Suarez-Farinas M, Li K, Fuentes-Duculan J, Hayden K, Brodmerkel C, Krueger JG. Expanding the psoriasis disease profile: interrogation of the skin and serum of patients with moderate-to-severe psoriasis. J Invest Dermatol 2012;132(11):2552-64.

Wong HY, Khosrotehrani K, Roy E. Whole-mount staining coupled to a UV-inducible basal cell carcinoma murine model. STAR Protoc 2021;2(1):100329.

Yao Y, Richman L, Morehouse C, de los Reyes M, Higgs BW, Boutrin A, et al. Type I interferon: potential therapeutic target for psoriasis? PLoS One 2008;3(7):e2737.

Zhou Y, Zhou B, Pache L, Chang M, Khodabakhshi AH, Tanaseichuk O, et al. Metascape provides a biologist-oriented resource for the analysis of systems-level datasets. Nat Commun 2019;10(1):1523.

### Figure S1

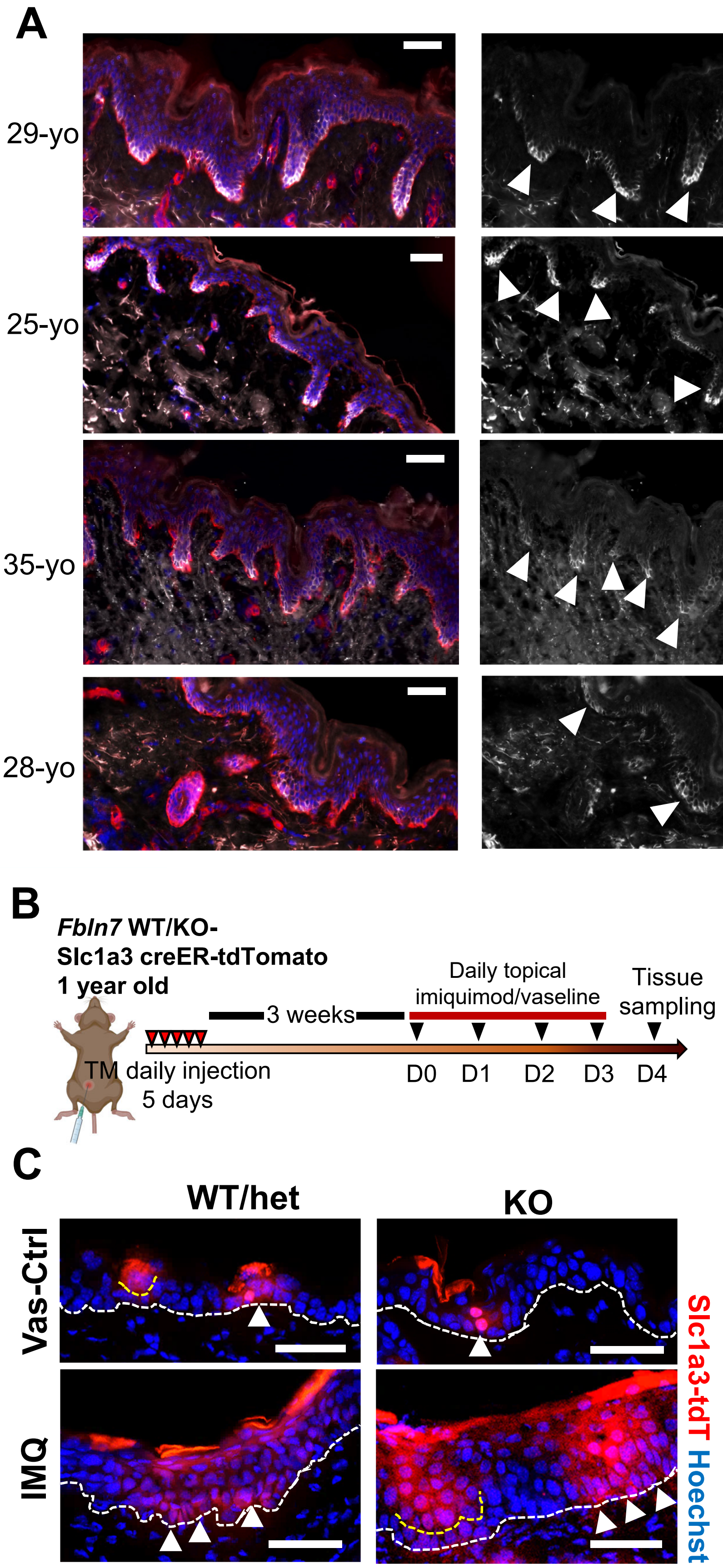

A
29-yo
25-yo
35-yo
28-yo
Slc1a3 Fibulin 7 Hoechst
B
Fbln7 WT/KO-
Slc1a3 creER-tdTomato
1 year old
3 weeks
Daily topical
imiquimod/vaseline
Tissue
sampling
TM daily injection
5 days
D0
D1
D2
D3
D4
C
WT/het
KO
Vas-Ctrl
IMQ
Slc1a3-tdT Hoechst

**Supplementary Figure 1. Fibulin 7 localization in human skin rete ridges proximal to Slc1a3-positive basal cells and genetic labelling of fast-cycling epidermal stem cells in the mouse model using *Slc1a3*-creER-tdTomato. (A)** Photomicrographs of normal abdominal human skin stained with anti-Slc1a3, fibulin 7 and Hoechst. Scale bars, 50 µm. White arrow heads indicate rete ridges. Yo, year old. **(B)** Tamoxifen (TM) was injected 5 times (once daily) into *Slc1a3*-creER-tdT mice in the background of *Fbln7* WT, heterozygous (het) or KO to activate fast-cycling stem cells labelling. 3 weeks later control or IMQ cream was applied 4 times (once daily) on the dorsal skin. **(C)** Td-Tomato labelled fast-cycling EpSCs in *Fbln7* WT or KO dorsal skin with/without IMQ-induced inflammation. Basal cells are epidermal cells at the dermal-epidermal junction (dashed white lines; examples are pointed by white arrow heads). Suprabasal cell clusters are ventrally demarcated with dashed yellow lines.

### Figure S2

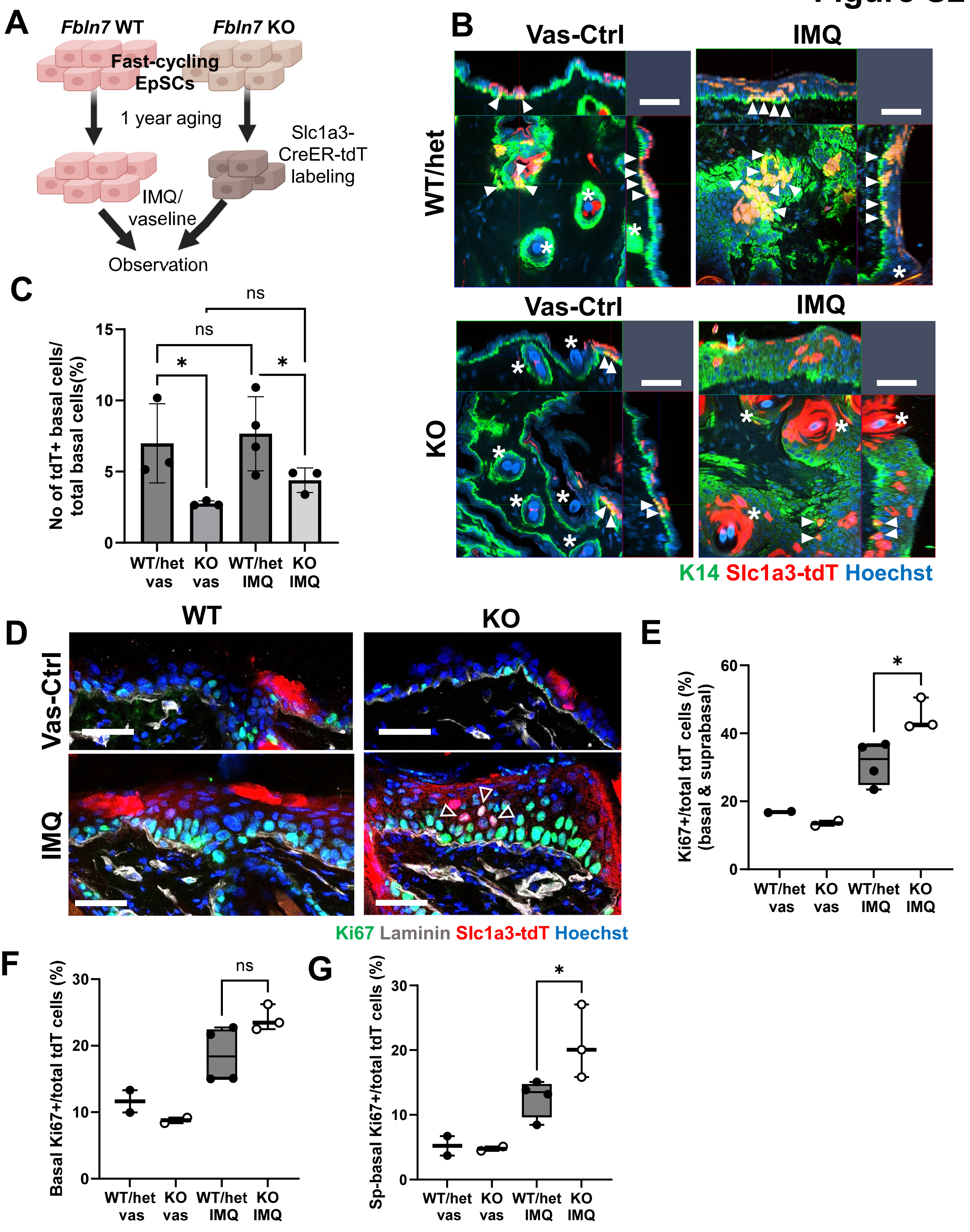

A
Fbln7 WT
Fbln7 KO
Fast-cycling EpSCs
1 year aging
Slc1a3-CreER-tdT labeling
IMQ/ vaseline
Observation
B
Vas-Ctrl
IMQ
WT/het
KO
K14 Slc1a3-tdT Hoechst
C
No of tdT+ basal cells/ total basal cells(%)
WT/het vas
KO vas
WT/het IMQ
KO IMQ
ns
*
D
WT
KO
Vas-Ctrl
IMQ
Ki67 Laminin Slc1a3-tdT Hoechst
E
Ki67+/total tdT cells (%) (basal & suprabasal)
F
Basal Ki67+/total tdT cells (%)
G
Sp-basal Ki67+/total tdT cells (%)

**Supplementary Figure 2. *Slc1a3*-expressing fast-cycling epidermal stem cells modulations in *Fbln7* WT/KO mice. (A)** Cartoon depicting fast-cycling EpSCs after 1-year aging at which point the stem cells are genetically labelled with fluorescent tdTomato (tdT) based on Slc1a3-creER inducible system, prior to Vas/IMQ treatment. **(B)** Z-stacked wholemount immunostaining images of WT/KO dorsal skin treated with vas-control or IMQ, timeline as figure 2C. K14, keratin 14 as epidermal basal cell marker. Examples of *Slc1a3*-tdT+ basal cells are marked with white arrow heads while hair follicles are marked with white asterisks. View from dermal side. **(C)** Graph summarizes *Slc1a3*-tdT+ basal cell counts from (B). Mean ± S.D. n = 3 for all groups except WT/het-IMQ, n = 4. Het, heterozygous. ANOVA with Tukey test; ns, not significant. * $P < 0.05$. All scale bars, 50 μm. **(D)** Immunofluorescence micrographs describing tdT-positive cells within the epidermis in control or IMQ-treated WT vs KO skin. Laminin staining was used to localize the BM. Cells that are double positive for Ki67 and tdT were quantified. White arrow heads indicate some tdT+ Ki67+ cells in the suprabasal layer. that are Scale bars, 50 μm. Hoechst, nuclear staining. **(E, F, G)** Graphs summarize cell count results from overall Ki67+ tdT-labelled cells within the interfollicular epidermis (IFE, excluding hair follicle cells), IFE basal cells only and IFE suprabasal (Sp-basal) cells only. n = 2 for WT or KO control-treated groups, n = 4 for WT/het (heterozygous) IMQ-treated group and n = 3 for KO-IMQ treated group. Unpaired T-tests. * $P < 0.05$, ns; not significant.

### Figure S3

**A**

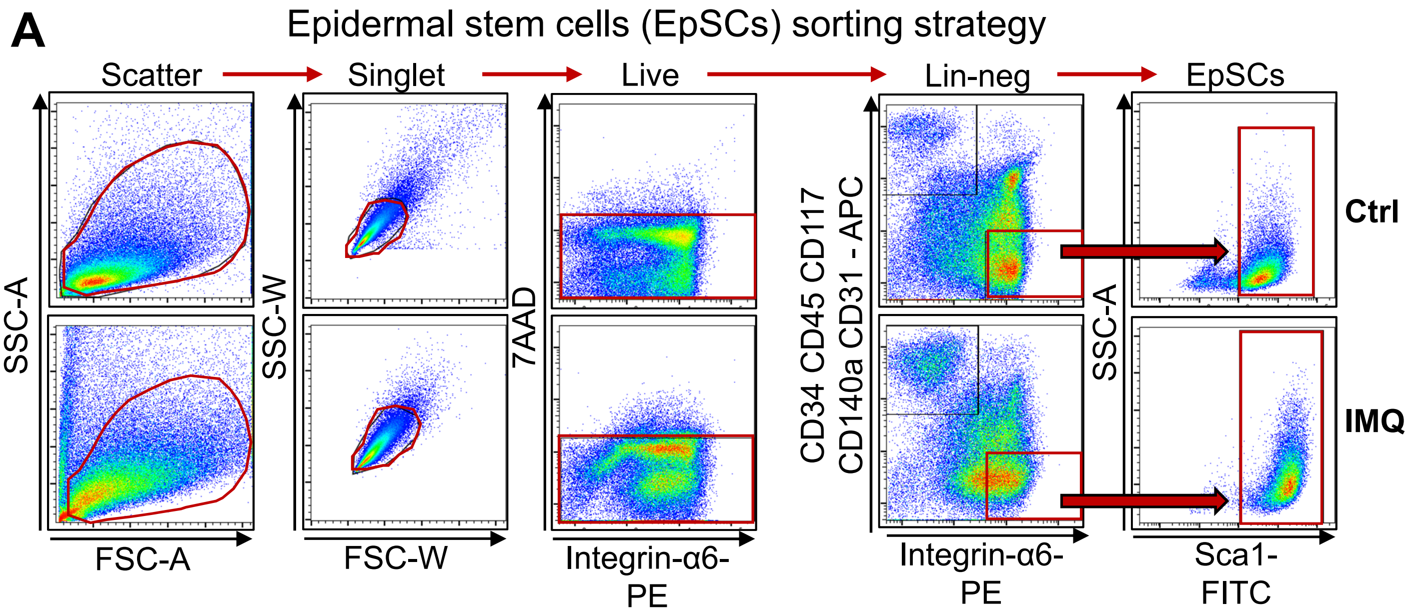


**B** **DEGs Vas-Ctrl vs IMQ (FC ≥ 2; FDR $P \leq 0.05$)**

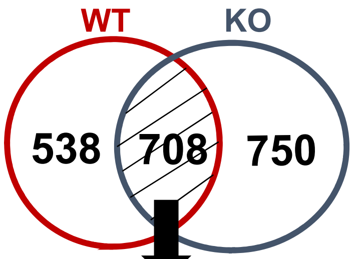


**C**

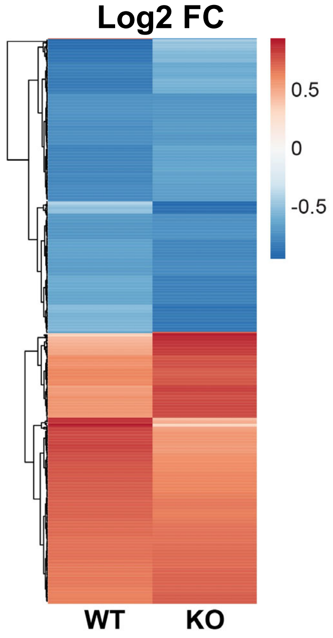


**D**

| Downregulated genes ontology | LogP |
|---|---|
| skin development | -13.5654 |
| triglyceride metabolic process | -7.88938 |
| extracellular matrix organization | -6.78252 |
| negative regulation of locomotion | -5.32014 |
| intermediate filament cytoskeleton organization | -4.48533 |
| negative regulation of angiogenesis | -3.78588 |

**E**

| Upregulated genes ontology | LogP |
|---|---|
| keratinization | -20.5243 |
| Microtubule-dependent trafficking from Golgi to the plasma membrane | -8.72528 |
| inflammatory response | -5.9522 |
| chemotaxis | -5.73569 |
| cell-cell adhesion | -5.66282 |
| EGFR interacts with phospholipase C-gamma | -5.19358 |
| glucose catabolic process | -5.18266 |
| response to lipopolysaccharide | -4.98372 |
| HIF-1 signaling pathway | -4.88741 |
| IL-17 signaling pathway | -4.8512 |
| vasculature development | -4.6345 |
| ERBB signaling pathway | -4.59087 |
| monocarboxylic acid transport | -4.47591 |
| cellular response to interleukin-1 | -3.48284 |

**Supplementary Figure 3. Basal epidermal stem cells sorting strategy and the common genes regulation post-imiquimod treatment. (A)** Flow cytometry diagrams, for sorting of EpSCs from the dorsal skin treated with vas-control or IMQ, indicating selection of live cells (7AAD negative) highly expressing both Integrin-α6 and Sca1 (basal EpSCs markers) while excluding other cell types (Lin-neg, lineage negative for CD34, CD45, CD117, CD140a, or CD31). **(B)** Venn diagram displays shared genes regulation between *Fbln7* WT and KO EpSCs after IMQ treatment for 4 days compared to control (shaded area, 708 genes), with **(C)** the heat map representation (log2 FC, fold change). **(D, E)** GO from genes down- and upregulated after IMQ treatment, respectively.

### Figure S4

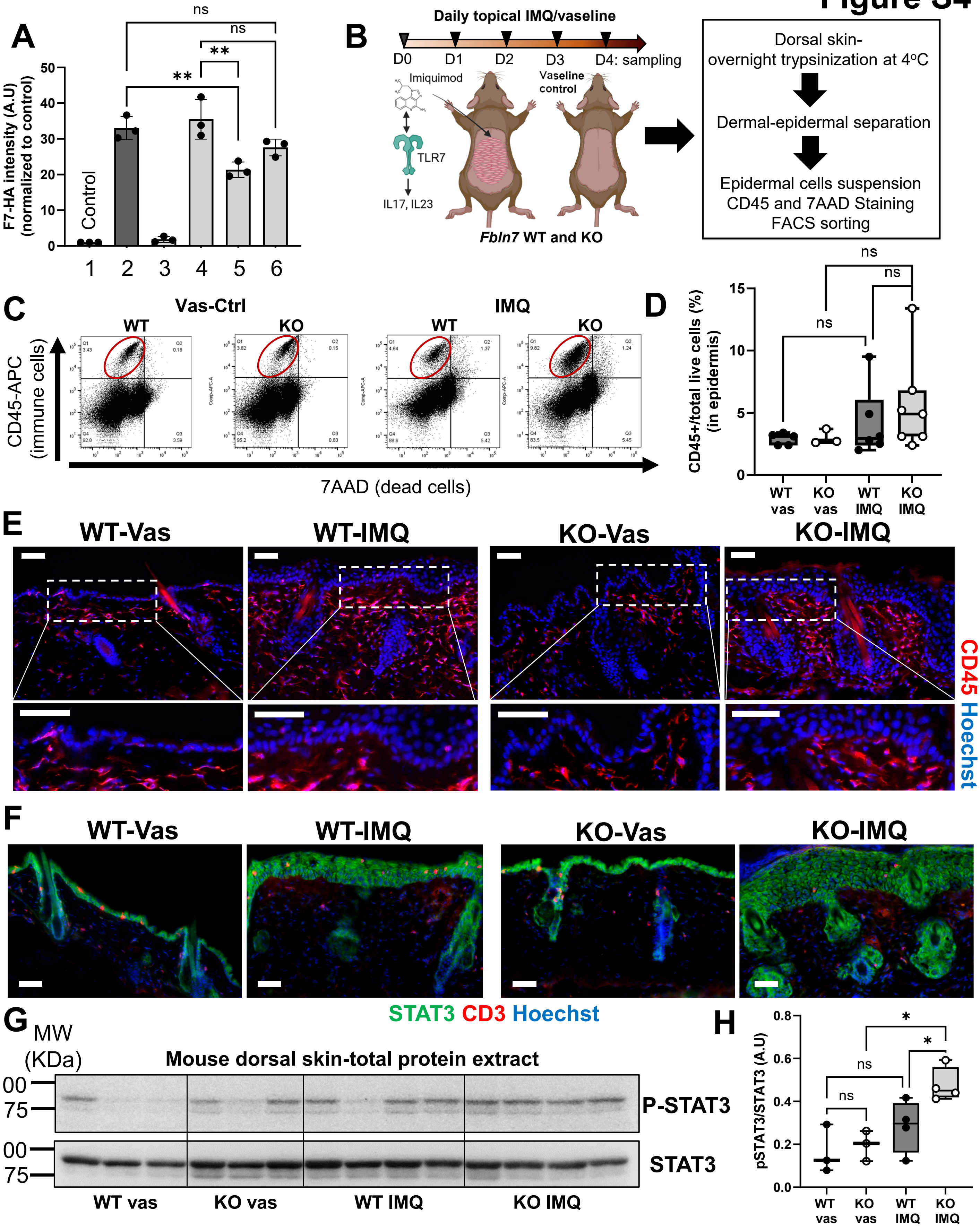

A
F7-HA intensity (A.U) (normalized to control)
Control
1 2 3 4 5 6
ns
**
B
Daily topical IMQ/vaseline
D0 D1 D2 D3 D4: sampling
Imiquimod
TLR7
IL17, IL23
Vaseline control
Fbln7 WT and KO
Dorsal skin- overnight trypsinization at 4°C
Dermal-epidermal separation
Epidermal cells suspension CD45 and 7AAD Staining FACS sorting
C
Vas-Ctrl
IMQ
WT
KO
CD45-APC (immune cells)
7AAD (dead cells)
D
CD45+/total live cells (%) (in epidermis)
WT vas
KO vas
WT IMQ
KO IMQ
E
WT-Vas
WT-IMQ
KO-Vas
KO-IMQ
CD45 Hoechst
F
STAT3 CD3 Hoechst
G
MW (KDa)
Mouse dorsal skin-total protein extract
P-STAT3
STAT3
WT vas
KO vas
WT IMQ
KO IMQ
H
pSTAT3/STAT3 (A.U)
*

**Supplementary Figure 4. Assessment of the number of epidermal immune cells in *Fbln7* WT or KO dorsal skin after imiquimod-induced inflammation. (A)** Immunoblot quantification from the same experiment shown in figure 4E (fibulin 7-HA blot), expressed as fold change to control. Bar graphs show mean ± S.D. ** $P < 0.01$, ns; not significant. **(B)** Schematics of experimental timeline for vas-control or IMQ administration in the skin and procedures taken before FACS analysis. **(C)** FACS charts designate the proportion of cells identified as live immune cells (CD45+/7AAD-) summarized in **(D)** graph from the overall data, n = 5 (WT-control), n = 3 (KO-control), n = 6 (WT-IMQ treated) and n = 7 (KO-IMQ treated). **(E, F)** Representative immunostaining micrographs of CD45+ total immune cells (e) and CD3+ pan-T cells/Stat3+ epidermal cells (f), 4 days after vas or IMQ application. White-dashed box areas were enlarged in the lower panels of (e). N=4 per group in CD45 staining; n = 4 and n = 6 in each WT or KO vas-control groups and WT or KO IMQ groups, respectively. Hoechst, nuclear staining. Scale bars, 50 μm. **(G, H)** Immunoblots display the abundance of STAT3 and phospho-STAT3 (Y705) from mouse dorsal skin extract (G) and its quantification (H). n = 3 and n = 4 for vas-control and IMQ-induced WT/KO mice, respectively. Statistical test used for all graphs: 2-way ANOVA, Tukey test. ns, not significant. * $P < 0.05$.

Figure S5

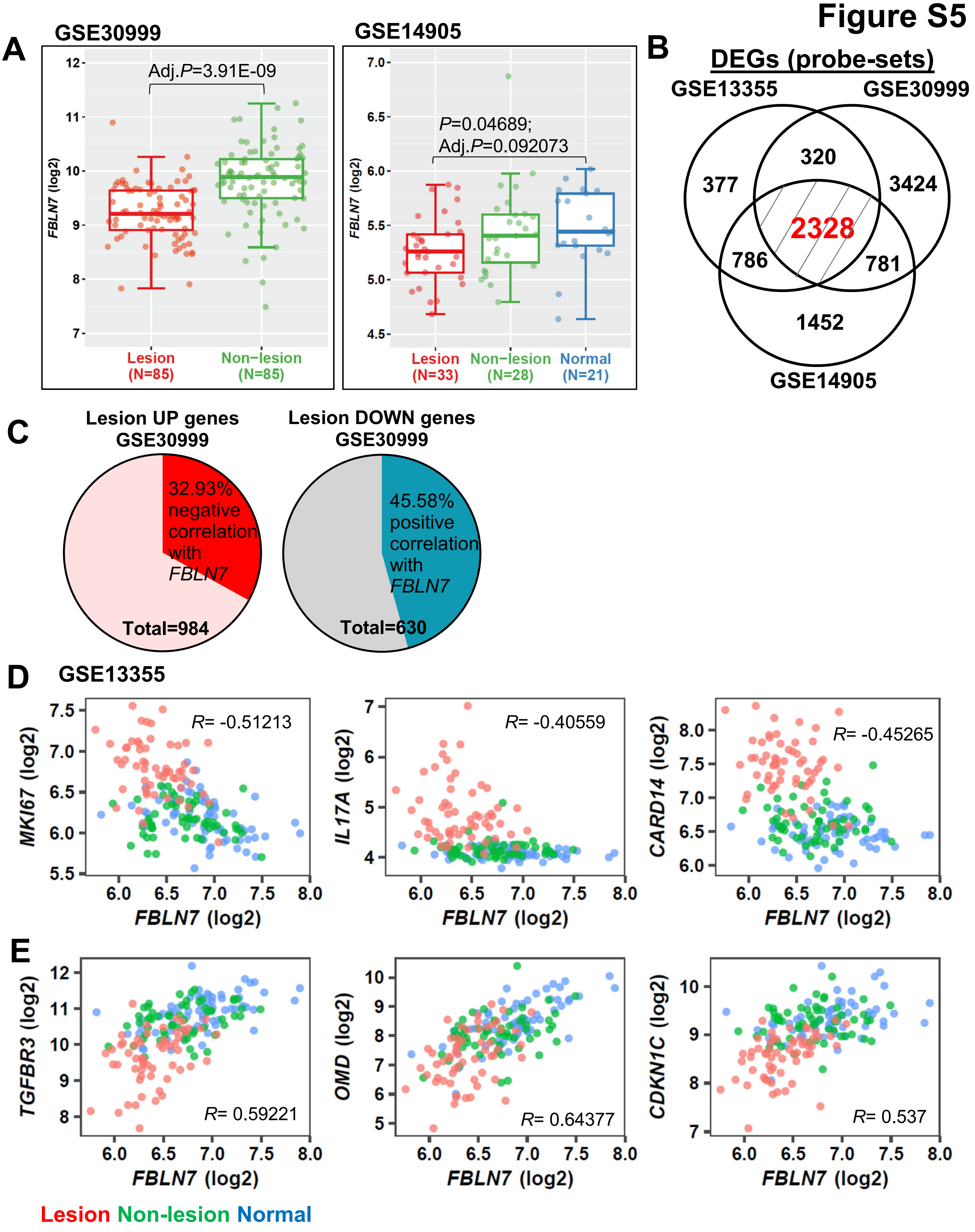

A
GSE30999
Adj.P=3.91E-09
FBLN7 (log2)
Lesion (N=85)
Non-lesion (N=85)
GSE14905
P=0.04689;
Adj.P=0.092073
Lesion (N=33)
Non-lesion (N=28)
Normal (N=21)
B
DEGs (probe-sets)
GSE13355
GSE30999
GSE14905
377
320
3424
2328
786
781
1452
C
Lesion UP genes GSE30999
32.93% negative correlation with FBLN7
Total=984
Lesion DOWN genes GSE30999
45.58% positive correlation with FBLN7
Total=630
D
GSE13355
MKI67 (log2)
R= -0.51213
IL17A (log2)
R= -0.40559
CARD14 (log2)
R= -0.45265
FBLN7 (log2)
E
TGFBR3 (log2)
R= 0.59221
OMD (log2)
R= 0.64377
CDKN1C (log2)
R= 0.537
FBLN7 (log2)
Lesion Non-lesion Normal

**Supplementary Figure 5. *FBLN7* expression and gene correlations from independent human psoriasis public datasets.** **(A)** *FBLN7* mRNA comparisons from GSE30999 and GSE14905. Each dot represents overall expression from one individual. **(B)** 2328 common DEGs shared between 3 datasets (data as gene_probe-sets) which translates to 1614 genes. **(C)** Proportions of *FBLN7* correlations with genes that are up or downregulated in psoriasis lesion from GSE30999. **(D)** Examples of *FBLN7* negative and **(E)** positive correlations with common psoriasis-related genes in GSE13355.

### Figure S6

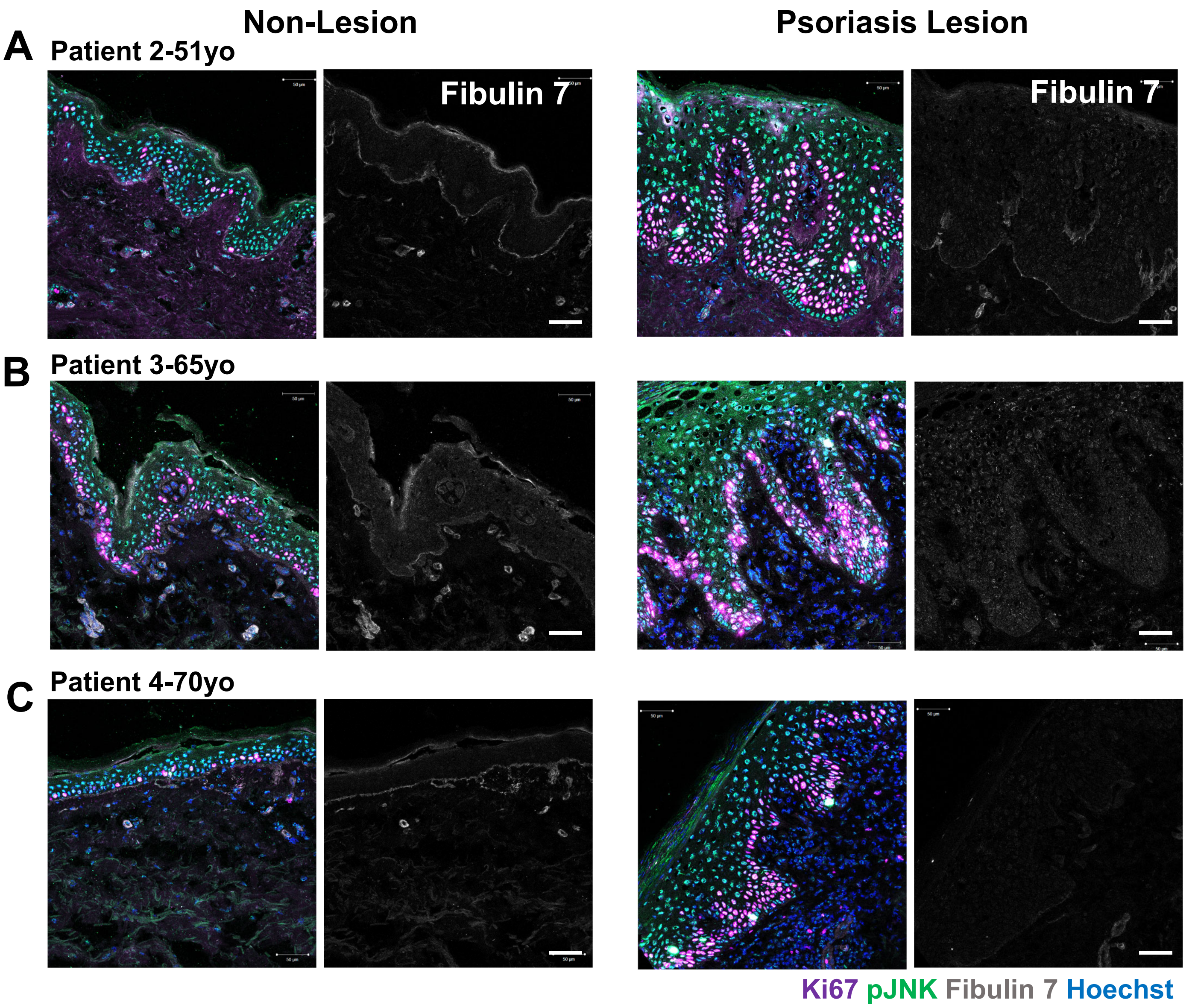


**Supplementary Figure 6. Immunostainings of fibulin 7 in human psoriasis lesion compared to non-lesion areas.** Photomicrographs from a 51-year-old **(A)**, 65-year-old **(B)** and 70-year-old skin sections. Scale bars, 50 µm.

**Supplementary Table 1.** DEGs between *Fbln7* WT and KO epidermal stem cells post-IMQ treatment.

**Supplementary Table 2.** List of genes that are more highly regulated upon loss of *Fbln7* in epidermal stem cells after IMQ treatment compared to control.

**Supplementary Table 3.** List of commonly regulated genes in *Fbln7* WT or KO epidermal stem cells after IMQ treatment compared to control.

**Supplementary Table 4.** List of genes that are more strongly affected in the WT mice after IMQ compared to control.

**Supplementary Table 5.** Commonly regulated genes in human psoriasis and their correlations to *FBLN7*. Data summarized from GSE13355, GSE30999, GSE14905.
